# Polycaprolactone-based shape memory foams as self-fitting vaginal stents

**DOI:** 10.1101/2024.01.26.577474

**Authors:** Ashley Hicks, Courteney Roberts, Andrew Robinson, Kailey Wilson, Varsha Kotamreddy, Trace LaRue, Arian Veyssi, Felipe Beltran, Julie Hakim, Manuel Rausch, Melissa Grunlan, Elizabeth Cosgriff-Hernandez

**Affiliations:** Department of Biomedical Engineering, The University of Texas at Austin, Austin, Texas, 78712; Department of Biomedical Engineering, Texas A&M University, College Station, Texas, 77843; Department of Aerospace Engineering & Engineering Mechanics, The University of Texas at Austin, Austin, Texas, 78712; Department of Obstetrics and Gynecology, Baylor College of Medicine, Houston, Texas, 77030

**Keywords:** Emulsion templating, shape memory polymers, vaginal stenosis

## Abstract

There is an urgent critical need for a patient-forward vaginal stent that can prevent debilitating vaginal stenosis that occurs in up to 75% of patients who undergo pelvic radiation treatments and adolescent patients after vaginal reconstruction. To this end, we developed a self-fitting vaginal stent based on a shape-memory polymer (SMP) foam that can assume a secondary, compressed shape for ease of deployment. Upon insertion, the change in temperature and hydration initiates foam expansion to shape fit to the individual patient and restore the lumen of the stent to allow egress of vaginal secretions. To achieve rapid actuation at physiological temperature, we investigated the effect of architecture of two photocurable, polycaprolactone (PCL) macromers. Star-PCL-tetraacrylate displayed reduced melting temperature in the target range as compared to the linear-PCL-diacrylate. Emulsion-templating was then used to fabricate foams from 75:25 water-in-oil (W/O) emulsions that were subsequently annealed to yield high-porosity SMP foams. Upon axial shape memory testing, both foams displayed excellent shape fixity (90%); however, only the PCL *star-*foams displayed shape recovery (∼84%) at 37°C to its permanent shape. A custom mold and curing system was then used to fabricate PCL *star-* foams into hollow, cylindrical stents. The stent was crimped to its temporary insertion shape (50% reduction in diameter, OD ∼ 11 mm) with a custom radial crimper and displayed excellent shape fixity for deployment (> 95%) and shape recovery (∼ 100%). To screen vaginal stents, we developed a custom benchtop pelvic model that simulated vaginal anatomy, temperatures, and pressures with an associated computational model. A hysteroscope was used to visualize stent expansion and deformation via a scope port near the cervix of the benchtop model. A crimped SMP vaginal stent was deployed in the model and expanded to walls of the canal (∼70% increase in cross-sectional area) in less than 5 minutes after irrigation with warm water. The vaginal stent demonstrated retention of vaginal caliber with less than 1% decrease in cross-sectional area under physiological pressure. Collectively, this work demonstrates the potential for SMP foams as self-fitting vaginal stents to prevent stenosis. Additionally, this work provides new open-source tools for the iterative design of other gynecological devices.

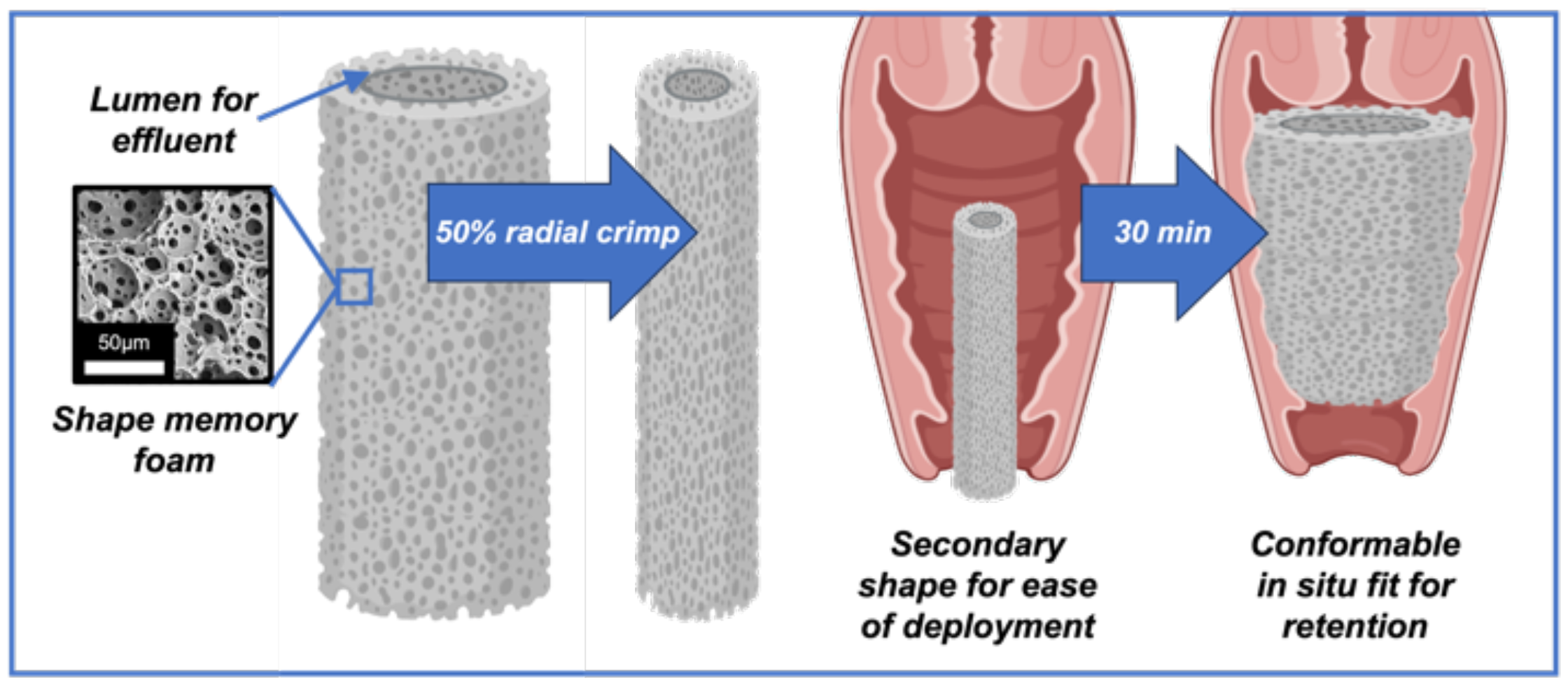

Created in BioRender

## 1. Introduction

There is an urgent clinical need for a vaginal stent that can address vaginal stenosis in post-surgical and post-radiation settings, impacting over 340,000 patients in North America each year.^1, 2^ The urgency in addressing vaginal stenosis is underscored by the shortcomings of current treatment options. Existing commercially available stents, including stainless steel variants and inflatable balloon stents, exhibit notable challenges with comfort and retention in the vagina and often require additional fixation to maintain their position in the canal. Stent expulsion can occur during simple routine motions like walking or sitting, making them highly ineffective solutions for most patients. As a result, physicians are left to craft makeshift stents from common medical equipment such as foley catheters, condoms, and medical foam.^3, 4^ This is particularly true in patients whose size is incompatible with current commercially available options, most often pediatric patients. The search for an alternative solution remains underdeveloped, resulting in ∼50% rate of secondary revision surgery for pediatric patients.^5^ The failure modes of these devices indicate that the design of a novel vaginal stent must rely on the premise of a conformal, retained fit within the vaginal cavity.

Shape memory polymers (SMP) are a class of material that present a unique opportunity to address this form-fitting requirement. SMPs are dual-shaped polymeric materials that can alter their geometry in a predefined manner in response to the appropriate environmental stimuli (e.g. heat, light, etc.).^6-8^ The permanent shape is determined by the initial fabrication step, whereas the temporary “fixed” shape is determined by tailored programming of the material. In comparison to shape memory alloys, SMPs enable greater tailoring of material properties (i.e. transition temperature, Young’s modulus, maximum elongation) to better meet functional engineering requirements. SMPs have been implemented in a broad range of form-fitting applications, historically dominated by the use of phase-segregated polyurethanes.^8^ Although polyurethanes are widely favorable for their long-term stability and highly tunable nature, their innate biostability is unfavorable for use in temporary implants due to the potential for residual wear particles to incite a chronic host response.^9, 10^ Polyesters provide an accommodative solution for this, as their hydrolytically labile ester bonds enable resorption and clearance of the material through the body’s immune response.^11-13^ Polycaprolactone (PCL) has gained notable prevalence in biomaterial applications such as stents, sutures, and bone grafts due to its established biocompatibility, tunable mechanical properties, and thermoresponsive shape memory capabilities.^13-23^ Its melt transition temperature (*T*_*m*_ ∼ 55°C) makes it a viable candidate for eliciting shape memory behavior without significant risk of thermal damage to surrounding tissues. Additionally, it is susceptible to hydrolytic degradation due to the presence of ester bonds in its backbone, making it a viable option for temporary implants. Thus, we selected to pursue the design of a self-fitting vaginal stent from PCL towards improving the available preventative measures for vaginal stenosis.

The vagina is a trophic, viscoelastic organ comprising multiple tissue layers and is acted upon by forces emerging from within the organ lumen and pelvis. The development of an effective vaginal stent necessitates robust testing modalities to evaluate its performance. While *in vivo* models offer valuable insights, they are not reasonable options for the preliminary design stages of a novel stent. Despite a recent initiative in the medical device field to design alternative models (*in vitro* or *in silico*) for design iteration, accessible models for vaginal conditions remain largely nonexistent. To address this, we will utilize clinical data to construct a testing apparatus for the iterative design of our SMP vaginal stent to evaluate the performance of the device under physiological conditions. Additionally, a computational model will be proposed as a supplemental tool, providing a more comprehensive understanding of vaginal mechanics. Establishing both benchtop and computational models requires dedicated resources, emphasizing the need for collaborative efforts to construct and validate these essential tools. Here, we will utilize an emulsion templating approach to fabricate an SMP foam from photocurable PCL macromers to achieve a self-fitting vaginal stent. Our proposed vaginal stent design assumes a secondary, compressed shape to facilitate insertion into the narrow vaginal introitus. Upon insertion, the change in temperature and hydration initiates foam expansion to shape fit to the individual patient. We hypothesize that the conformal fit will anatomically secure the stent in place to prevent egress, prevent tissue apposition and provide tissue stretch during healing to limit fibrosis, and restore the lumen of the stent to allow egress of vaginal secretions. To achieve target self-fitting and mechanical properties of PCL-based foams, we investigated effects of polymer chemistry and emulsion variables on deployment and retention outcomes. First, we investigated two polymer architectures, *linear*-PCL-diacrylate (DA) and *star*-PCL-tetraacrylate (TA), to identify an SMP foam with rapid shape recovery at body temperature. The effect of internal phase volume of the emulsion on shape recovery and mechanical properties was investigated to balance lumen retention and patient comfort. To expedite stent screening and provide better predictions of *in vivo* performance, a custom benchtop pelvic model was developed to simulate vaginal anatomy, temperatures, and pressures. We used this benchtop apparatus to test deployment, shape recovery, and lumen deformation under physiological pressures. Overall, we aim to demonstrate the potential of this newly designed SMP foam for use as a self-fitting vaginal stent and provide novel *in vitro* and *in silico* pelvic models for guiding the design of future gynecological devices.

## 2. Materials & Methods

### 2.1 Materials

*Linear*-PCL-diol (M_n_ = 10k gmol^-1^ per manufacturer specifications), ε-caprolactone, pentaerythritol, tin(II) 2-ethyl hexanoate (Sn(Oct), methanol, dichloromethane (DCM), 4-(dimethylamino)pyridine (DMAP), triethylamine (Et_3_N), acryloyl chloride, ethyl acetate, potassium carbonate (K_2_CO_3_), anhydrous magnesium sulfate (MgSO_4_), and deuterated chloroform (CDCl_3_). All solvents were dried over 4 Å molecular sieves, all reagents were vacuum-dried overnight, and all glassware and stir bars were dried at 120 °C prior to use. Polyglycerol polyricinoleate 4125 (PGPR) was donated by Paalsgard. All other chemicals were purchased from Sigma Aldrich.

### 2.2 Macromer Synthesis

Reactions were run under a nitrogen (N_2_) atmosphere with a Teflon-covered stir bar. Following purification, molecular structures (including % acrylation, architecture, and *M*_*n*_) were confirmed with ^1^H NMR spectroscopy (Avance Neo 400 MHz spectrometer) with CDCl_3_ as the standard. The target *M*_*n*_ and architecture of *linear*-PCL-DA and *star-*PCL-TA macromers were verified by comparing the repeat unit C***H***_**2**_ *d* = 4.1 ppm versus the terminal C***H***_**2**_ *d* = 3.7 ppm.

*Linear*-PCL-diol (10k gmol^-1^) was purchased from Millipore Sigma while *star*-PCL-tetrol (10k gmol^-1^) was synthesized via ring opening polymerization (ROP) per previously established protocols where ε-caprolactone (20.0 g), a tetrafunctional initiator (pentaerythritol), and Sn(Oct)_2_ were combined in a round bottom (rb) flask and maintained at 120°C overnight.^23^ The targeted M_n_ (10k gmol^-1^) was achieved by maintaining the monomer-to-initiator molar ratio (88:1, [M]:[I]). The crude product was then purified and vacuum dried overnight (RT, 30 in. Hg).

*Linear****-***PCL-diol and *star-*PCL-tetrol were each acrylated to form photocrosslinkable *linear*-PCL-DA and *star-*PCL-TA macromers, respectively. Each diol or tetrol (20.0 g) was combined with DMAP (6.6 mg fo**r** *linear****-***PCL-diol; 13.2 mg for *star-*PCL-tetrol) and dissolved in DCM (120 mL). Following purging with N_2_, Et_3_N and acryloyl chloride (4.0 mmol & 8.0 mmol for *linear***-**PCL-diols; 8.0 mmol & 16.0 mmol for *star*-PCL-tetrol) were each added dropwise to the flask and stirred under positive pressure for 30 min at RT. Established work-up procedures were followed to obtain *linear*-PCL-DA and *star*-PCL-TA.^23, 24^ The final products were isolated and vacuum dried overnight (RT, 30 in. Hg).

### 2.3 PolyHIPE Fabrication

PolyHIPE foam specimens were fabricated from photoinitiated PCL macromers (**Table S1**) with 75:25 water-in-oil (W/O) emulsions and subsequently annealed to yield highly porous shape memory foams (**Figure 1**). Briefly, macromers were individually combined with N-vinylpyrrolidone (NVP) and dissolved in toluene (30:5:65% w/w, respectively) to serve as the continuous phase of the emulsion. The photoinitiator phenylbis(2,4,6-trimethylbenzoyl) phosphine oxide (BAPO; 2.5% w/w) and PGPR surfactant (2.5% w/w) were added to the continuous phase and spun at 2500 RPM using a Flaktek Speedmixer for 2.5 minutes to form a homogenous solution. Aqueous calcium chloride solution (1% w/v) was added to the organic phase in six additions (75% v/v) and mixed at 500 RPM for 2.5 minutes per addition. Once a stable emulsion was formed, as confirmed visually, HIPEs were transferred to 3mL syringes (*d* ∼ 8mm). using the plunger to remove any trapped air. SMP foams were photocured for 4 hours on high (Analytik Jenka UV Transilluminator), rotating syringes 180° at 10, 20, 70 min, 2, and 3 hrs. After curing, specimens were removed from their mold and vacuum dried overnight to remove the diluent and internal phase. Specimens were annealed in 85°C DI water for 1 hour and cooled for 24 hours *in vacuo* to yield the final shape memory foam specimens.

**Figure 1:**
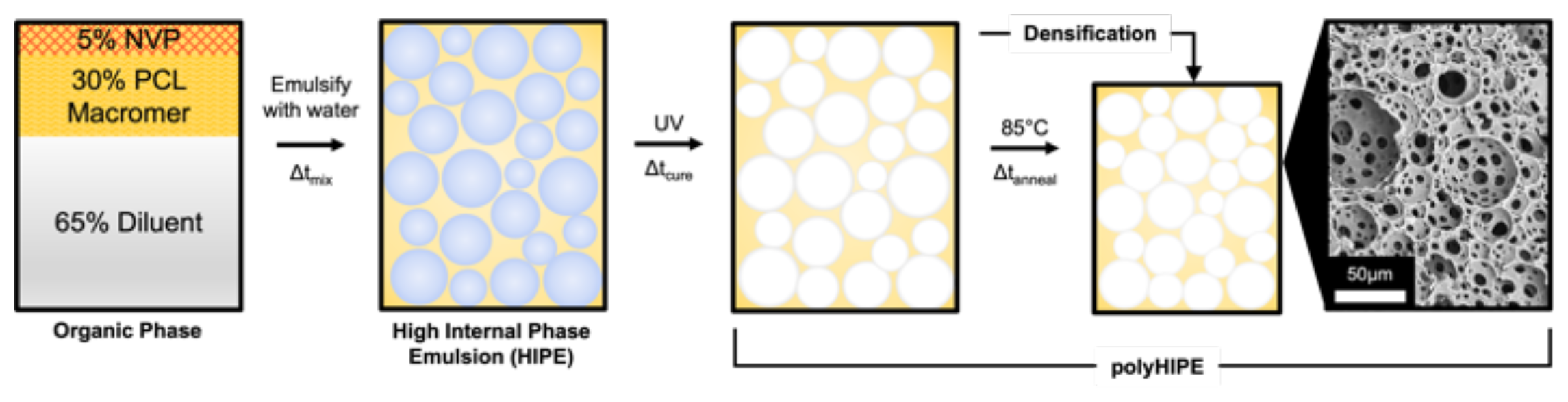
General scheme of emulsion templating of shape memory foams. The internal phase is gradually added into the organic phase while the system is mixed, forming a high internal phase emulsion (HIPE). The material is then cured to form a polymerized high internal phase emulsion (polyHIPE).

The effect of internal phase volume fraction on shape memory behavior and mechanical properties was also determined using *star*-PCL-TA polyHIPEs. Specimens were prepared as described above with three volume fractions (70%, 75%, and 80% v/v).

### 2.4 Gel Fraction

The gel fraction was measured gravimetrically to evaluate the extent of network formation. First, samples (*d* ∼ 6 mm, *h* ∼ 1 mm, n = 5) were weighed after drying *in vacuo* for 48 hours. The samples were placed in dichloromethane (DCM) for 4 hours at 1 mL / 10 mg of the sample, and vacuum dried again until a constant mass was achieved. The final weight divided by the initial weight was assessed as the gel fraction and corrected for the mass of the surfactant that would dissolve in the DCM.

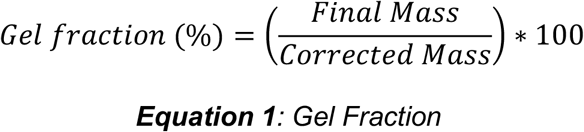

### 2.5 Porosity

The porosity of PCL polyHIPEs was determined gravimetrically. First, the polymer density was measured from solid film specimens of each macromer (*d* ∼ 6 mm, *h* ∼ 2 mm, n = 5). Dried HIPE samples (*d* ∼ 6 mm, *h* ∼ 2 mm, n = 5) were measured with electronic calipers and weighed. Porosity was calculated as a percent difference in the density of foams and analogous films.

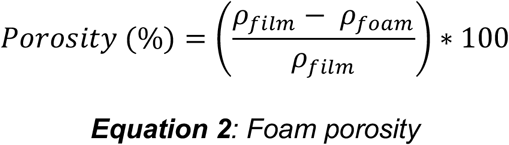

### 2.6 Pore Size

PolyHIPE samples (*d* ∼ 6 mm, *h* ∼ 10 mm, *N* = 3) were vacuum dried for 24 hours to remove the water before the characterization of the pore architecture. Scanning electron microscopy (SEM, Phenom Pro, Nanoscience Instruments) was used to characterize the average pore size for each composition. Samples were flash-frozen in liquid nitrogen to preserve pore architecture and sectioned into ∼ 1 mm thick pieces with a razor blade. Specimens were sputter-coated with gold with a thickness of ∼5 nm and imaged to yield nine images. Pore size measurements were conducted by measuring the diameters of the first ten pores that crossed the midline of each 1000x magnification micrograph. Average pore sizes for each polyHIPE composition are reported. A statistical correction was calculated to account for the non-perfect spherical pores, *h*^*2*^=*R*^*2*^-*r*^*2*^, where R is the void diameter’s equatorial value, r is the diameter measured from the micrograph, and h is the distance from the center of the pore.^25^ The average pore size values were multiplied by this correction factor, resulting in a more accurate description of the pore diameter.

### 2.7 Thermal Properties

The melting temperature (*T*_*m*_) and respective onset (*T*_*m,onset*_) of polyHIPEs were determined by differential scanning calorimetry (TA Instruments, DSC 250). Samples (*m* ∼ 5 mg, *n* = 3) were sealed in hermetic pans and heated from 0 to 200°C at a rate of 5 °C min^-1^. Values were taken from a second DSC cycle to eliminate thermal history. The onset and midpoint of PCL crystalline lamellae were measured using TA Universal Analysis software at the onset and maximum of the endothermic melt peak, respectively.

### 2.8 Axial Shape Memory

Shape fixity (*R*_*f*_) and shape recovery (*R*_*r*_) of polyHIPE specimens were first assessed using a cylindrical foam geometry (*d* ∼ 6 mm, *h* ∼ 10 mm, n = 4). First, a crimping temperature (*T*_*crimp*_) was determined to be the onset of the melt transition temperature of PCL (*T*_*m,onset*_; *linear*-PCL-DA = 50°C, *star*-PCL-TA = 40°C). A heating mantle equipped with a digital temperature probe was used to warm DI water to appropriate *T*_*crimp*_. Next, samples were subjected to the following protocol: (1) submerged into water bath previously heated to the designated *T*_*crimp*_ and maintained for 5 min; (2) removed and immediately crimped to 50% of the original length with a crescent wrench; (3) maintained in the wrench for 30 min to fix the new temporary shape in a 37°C incubator; (4) removed from the incubator and cooled in the wrench for 30 minutes, (5) removed from the wrench and allowed to sit for 30 min; (6) re-submerged into the water bath at 37°C for 5 min to elicit shape recovery, removed, allowed to cool at RT for 2 min. From this process, the shape fixity and shape recovery were quantified, where e_m_ is the maximum strain following step 2, e_u_ (N) is the strain in the stress-free state following step 4, and e_p_ is the final recovered strain following step 6. Strain values were determined via electronic caliper measurements. Two cycles (N) of shape memory testing were conducted.

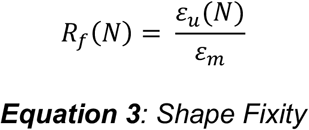

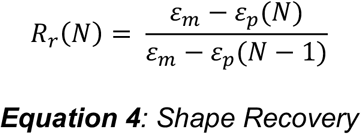

### 2.9 Compressive Properties

The polyHIPE compressive properties were tested with an Instron 3300, equipped with a 100-N load cell. The polyHIPEs were cured in 20 mL scintillation vials (t = 4hrs) for mechanical testing. Following annealing and densification, tops and bottoms of each disc specimen were removed with a razor blade to create flat parallel faces. Each specimen was sectioned into disks with a 3:1 diameter-to-height ratio to comply with ASTM standards, (*d* ∼ 15 mm, *h* ∼ 5 mm) and compressed at a strain rate of 0.05 mm/s. The compressive modulus was calculated from the slope of the linear region at 2% after correcting for zero strain. The moduli averages of the five disks were reported.

### 2.10 Degradation

The impact of hydrolysis on foam mechanical integrity was evaluated for both prepubescent (pH ∼7.4) and adult (pH ∼4.5) vaginal conditions. Disc specimens (h ∼ 5mm, d ∼ 15mm) were placed in 50mL solution (PBS or 150μM HCl) and incubated at 37°C for four weeks. Solutions were changed weekly to provide sink conditions, and dried foams were weighed to quantify mass loss at weekly time points. After the final timepoint, the compressive modulus was reassessed as previously described.

### 2.11 SMP Stent Fabrication

To fabricate stents from *star*-PCL-TA HIPEs, custom equipment was first designed and constructed. A dynamic curing system was implemented to allow for uniform photoinitiation by rotating the polyHIPE stent in its mold above the UV source (**Figure 2a**). The system also implements a UV strip light (Waveform Lighting, 365nm) to facilitate UV initiation within the stent lumen. Vaginal stents were fabricated from *star*-PCL-TA 75:25 polyHIPEs in the shape of a uniform hollow cylinder and processed as previously described (**Figure 2b**). Molds (*OD* ∼ 3.3 cm, *ID* ∼ 2.5 cm, *l* ∼ 152 cm) were created from concentric glass tubes and held in place with 3D-printed caps to ensure an even wall thickness along the length of the stent. Emulsion was loaded into molds using a 50mL syringe and silicone spacers were placed before adding the mold cap to fill the void space and remove trapped air bubbles. Stents were rotated at 20 RPM under UV illumination for 4 hours. The resulting polyHIPE stents were removed from their mold, dried, and annealed as previously described.

**Figure 2:**
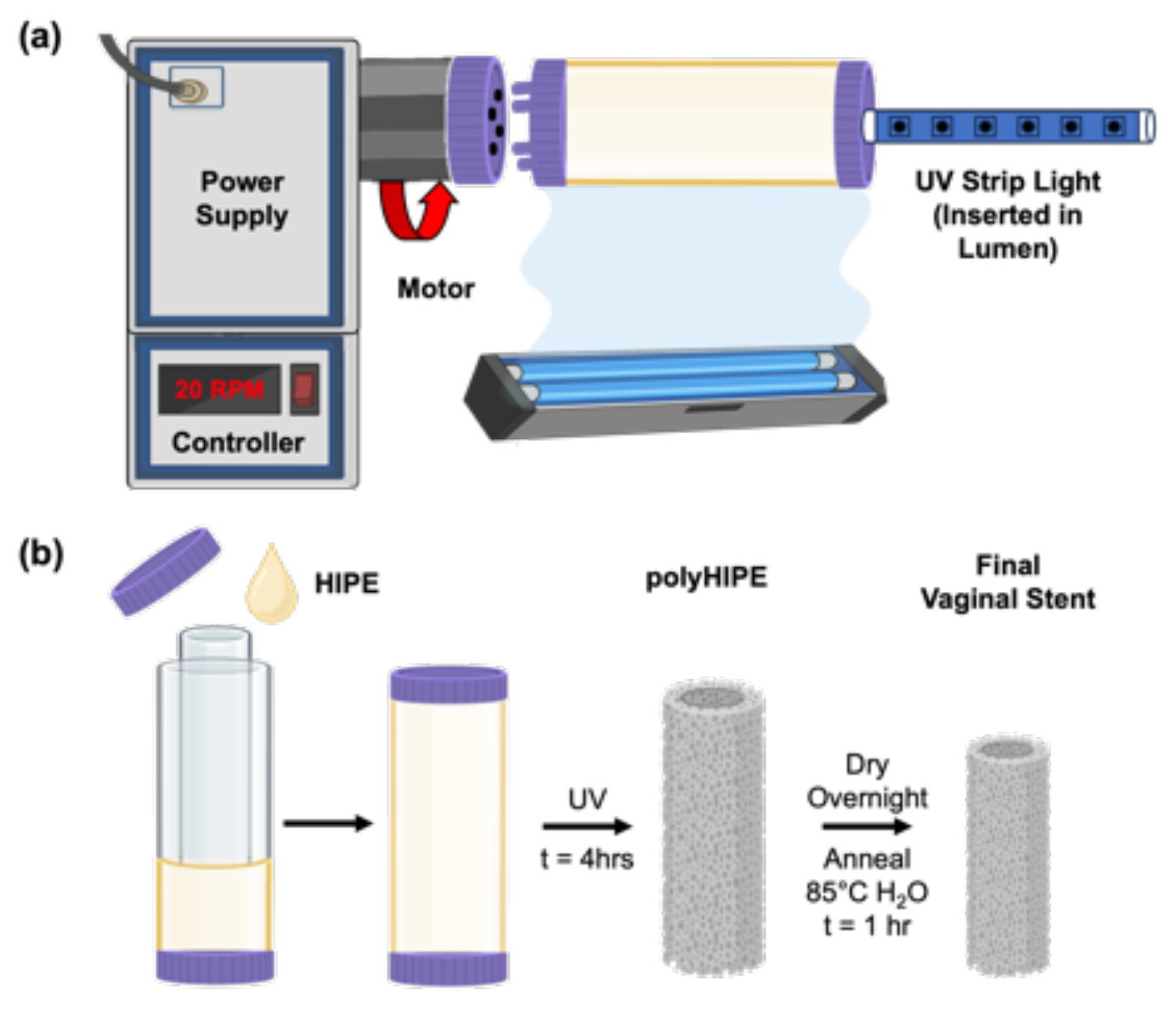
Fabrication of PCLTA polyHIPE vaginal stents. (a) Custom rotational device for UV curing of the vaginal stent. (b) General scheme of stent fabrication demonstrating mold setup and densification after fabrication. Created in BioRender.

### 2.12 Radial Shape Memory of Stents

Shape memory behavior of *star*-PCL-TA polyHIPE vaginal stents were assessed similarly to axial shape memory testing, as described above. Stents (*OD* ∼ 22 mm, *ID* ∼ 10 mm, h ∼ 7 cm, *N* = 3) were subjected to the following protocol: (1) submerged into water bath previously heated to the designated *T*_*crimp*_ = 40°C and maintained for 30 min; (2) removed and immediately crimped to 50% of the original length with a radial crimping device; (3) maintained in the crimper for 30 min to fix the new temporary shape; (4) removed from the crimper, placed in 5mL test tubes, and dried overnight; (5) removed from tubes and allow to rest for 30 minutes, (6) re-submerged into the water bath at 45°C for 5 min to elicit shape recovery, removed, and allowed to cool at RT for 2 min. From this process, the shape fixity and shape recovery were quantified, where e_m_ is the maximum strain following step 2, e_u_ (N) is the strain in the stress-free state following step 5, and e_p_ is the final recovered strain following step 6. Strain values were determined via electronic caliper measurements.

### 2.13 Benchtop Testing Apparatus Validation

A benchtop pelvic model was designed in collaboration with Lazarus 3D. Briefly, a vaginal canal was modeled from EcoFlex and enclosed in an acrylic pressure chamber. Heating tape (Briskheat, 144W/120V) was wrapped around the exterior of the acrylic chamber to heat the vaginal canal to physiological temperatures. Temperatures were measured in triplicate using a thermometer within three sections of the vaginal canal to ensure even heating: the cervical region, central region, and introitus region. An air pump was connected to the pressure chamber via silicone tubing and used to pressurize the canal to simulate resting pressures and pelvic floor muscle (PFM) contractions. A MizCure perineometer (OWOMED) was used to assess radial pressures within the model canal. The protocol for pressure assessment was adapted from clinical reports to make direct comparisons between the model and clinical pressure values.^26^ A hysteroscope (Endosee) was used to visualize stent expansion and deformation via a scope port near the cervix of the model.

### 2.14 Benchtop Stent Deployment & Retention

Prior to insertion, stents were subjected to 40°C for 30 minutes and radially crimped to 50% dimensions. The stent was maintained in the crimper for 30 minutes, removed and allowed to rest for 30 minutes before taking initial measurements. Following preparation, the crimped stent was inserted through the introitus of the model and immediately imaged to capture a reference for stent expansion. Stents were then irrigated with 100 mL of warm water (*T* ∼ 45°C) through the introitus and imaged to capture expansion. Stents were maintained in the canal overnight to equilibrate to physiological temperature. The model was then pressurized to simulate resting vaginal pressures and PFM contraction. Images were captured of stent deformation at each stage. Lumen deformation and stent egress from the model were both monitored throughout model pressurization to confirm successful retention.

### 2.15 Computational Modeling of Vaginal Anatomy & Stent Mechanics

A computational model of the benchtop system was developed. Briefly, the same computer-aided design model as used for the physical benchtop pelvic model was used for the computational pelvic geometry. To this end, the model was exported from SolidWorks (Version 2023, Dassault Systeme) as triangulated surfaces. The triangulated surfaces were then imported into the open-source meshing software TetGen (Version 1.6.0, WIAS Software). Next, the vaginal volume was meshed with 5598 linear tetrahedral elements and imported into the open-source nonlinear finite element software FEBio (Version 2.3.0, www.febio.org). In FEBio, we assigned the pelvic volume a 1-term Ogden, hyperelastic material model and informed its EcoFlex-specific material parameters.^27^ Additionally, we applied displacement and pressure boundary conditions to match those of the benchtop model.

### 2.16 Statistical Analysis

The data are displayed as mean ± standard deviation for each composition. To assess the significance of the data, we employed an analysis of variance (ANOVA) for multiple composition comparisons, supplemented by Tukey’s multiple comparisons test. Additionally, a Student’s t-test was conducted to ascertain any statistically notable distinctions between compositions. All tests were conducted at a 95% confidence interval (p<0.05).

## 3. Results & Discussion

### 3.1 Effect of Polymer Architecture on SMP PolyHIPE Properties

*Linear*-PCL-DA and *star*-PCL-TA successfully formed high internal phase emulsions (HIPEs) with a 75% internal phase as confirmed visually. Following annealing and densification of the polymer network (**Figure S2**), gel fraction studies revealed ∼90% incorporation of PCL macromers and small molecule crosslinker (NVP) in both polyHIPEs, indicating both HIPEs underwent effective crosslinking. Gravimetric porosity of *linear*-PCL-DA polyHIPEs demonstrate a ∼5% decrease from *star*-PCL-TA polyHIPEs (*linear* ∼ 65%, *star* ∼ 70%) when fabricated with identical internal phase volumes, likely attributed to the higher density of the *linear*-PCL-DA crystalline lamellae. The densification of the polymer network (occurs during removal of diluent and subsequently annealing) is dictated by the crystallization of the polymer chains. Linear polymer architectures possess an enhanced ability to pack together to form crystalline lamella in comparison to branched architectures, which are more sterically hindered.^28^

The crystalline lamellae of chemically crosslinked networks of PCL act as the switching segments of the shape memory system, allowing for temporary reshaping of the material as the crystalline lamellae melt and become malleable. Crystallization of the lamellae after reshaping the material permits shape fixity, or the ability of a material to maintain a secondary geometry. The chemical crosslinks serve as permanent netpoints, providing the network with the ability to “remember” its permanent shape upon remelting the crystalline regions. The branched architecture (**Figure 3a**) of *star*-PCL-TA in comparison to its *linear* counterpart partially impedes the ordered packing of crystalline regions thereby lowering its melt transition temperature. DSC thermograms of polyHIPE foams (**Figure 3b**) revealed a sharp, narrow peak at the melt transition of *linear*-PCL-DA (T ∼ 52°C). In comparison, the *star*-PCL-TA architecture demonstrates a broader, lower intensity melt peak (T ∼ 42°C) indicating a difference in crystallinity of the two polyHIPE foams. This information was used to determine an effective crimping (i.e. softening) temperature (*T*_*crimp*_) of the SMP foam for shape memory studies (*T*_*crimp,linear*_ = 50°C, *T*_*crimp,star*_ = 40°C).

**Figure 3:**
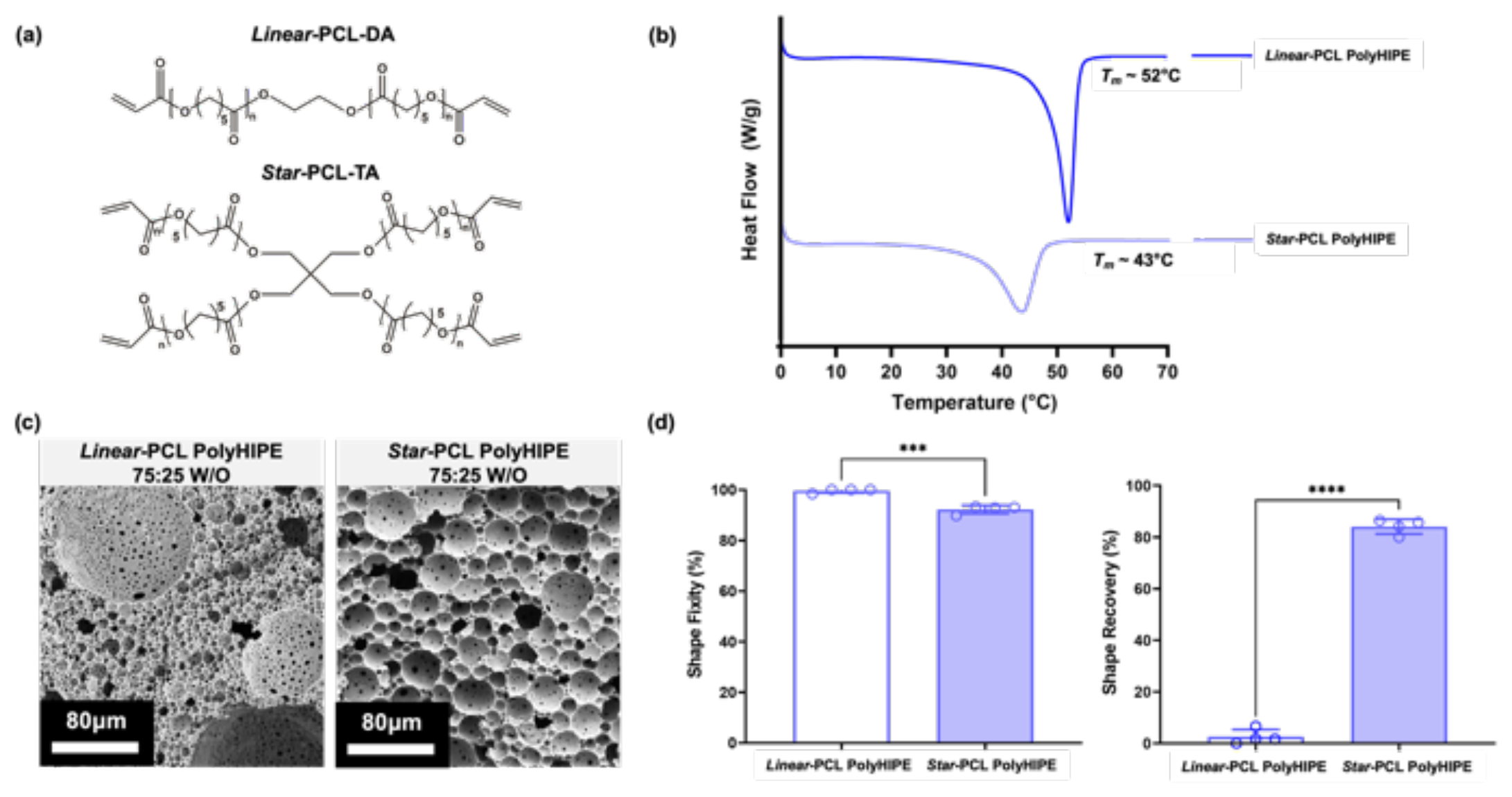
Structural properties of synthesized *linear*-PCL-DA and *star*-PCL-TA polyHIPE foams. (a) Schematic representing the chemical structures of the *linear*-PCL-DA and *star*-PCL-TA. (b) DSC thermograms of *linear* and *star* foam compositions demonstrating a decrease in *T*_*m*_ with *star* architecture. (c) Representative SEM images of *linear* and *star* foams. (d) Shape memory behavior of *linear* and *star* foams, including shape fixity and shape recovery values (T = 37°C, hydrated, n = 4). *** = p < 0.001, **** = p < 0.0001.

The microarchitecture of PCL polyHIPEs (**Figure 3c**) illustrate the difference in pore size distribution between the two polymer architectures (**Figure S3**). The presence of large pores (>100μm) within *linear*-PCL-DA foams is attributed to the higher viscosity of the *linear*-PCL-DA HIPE organic phase, partially impeding droplet breakup during emulsification. In comparison, the *star*-PCL-TA macromer solution is less viscous, allowing for uniform droplet dispersion and providing a narrower pore size distribution. The difference in macromer solution viscosity is a result of greater chain entanglement permitted by the *linear* polymer architecture.^29, 30^

The highly porous nature of the resulting PCL foams enables high compressibility (>50%) without fracturing, which is advantageous in the design of a self-expanding stent. Interconnects between the pores, formed as the pore walls begin thinning during densification, further enable compressibility by preventing air from being trapped within closed cells. A 50% crimp was integrated into the shape memory protocol to evaluate the effect of high strain on shape memory behavior (**Figure 3d**), which supports our design criteria for the self-fitting stent by permitting a crimped, temporary shape for ease of insertion into the narrow introitus of the vagina (∼1-4cm).^31^ Upon axial shape memory testing, both PCL macromers had excellent shape fixity (*R*_*f*_ > 90%) of the temporary crimped shape. However, the *linear*-PCL-DA polyHIPEs were not able to recover their permanent shape upon exposure to physiological temperatures. As anticipated, the higher ordered crystals within the *linear* network did not melt at 37°C, but the lower ordered crystals within the *star*-PCL-TA network enabled a recovery >85%. The recovery of the material can be further improved through temporary exposure to temperatures >37°C (e.g. irrigation with 45°C water), although it is important to consider the thermal sensitivity of relevant tissues. From this data, we hypothesize that the *star*-PCL-TA polyHIPE material will expand and conform to the vaginal canal through melting lower order crystals while maintaining its higher order crystallinity to provide structural rigidity. These findings support the potential of the *star*-PCL-TA polyHIPE to meet the success criteria outlined for the self-fitting vaginal stent.

**Table 1.** Structural properties of linear-PCL-DA and star-PCL-TA polyHIPEs (75:25 W/O).

| Macromer | Gel Fraction (%) | Porosity (%) | Pore Size ($\mu\text{m}$) | $T_{m, \text{onset}}$ ($^{\circ}\text{C}$) | $T_m$ ($^{\circ}\text{C}$) | Shape Fixity (%) | Shape Recovery (%) |
| --- | --- | --- | --- | --- | --- | --- | --- |
| *Linear*-PCL-DA | 88.3 ± 1.9 | 65.9 ± 1.4 | 32.7 ± 43.3 | 48.4 ± 0.0 | 52.0 ± 0.1 | 99.6 ± 0.8 | 2.1 ± 2.1 |
| *Star*-PCL-TA | 91.7 ± 0.7 | 70.5 ± 0.0 | 24.6 ± 13.6 | 36.4 ± 0.3 | 43.0 ± 0.5 | 92.3 ± 1.6 | 84.1 ± 2.9 |

### 3.2 Effect of Emulsion Internal Phase Volume on SMP PolyHIPE Properties

The effect of the SMP HIPE’s internal phase volume on resulting foam porosity was used to influence the *star*-PCL-TA polyHIPE’s mechanical properties. Based on the principles of emulsion templating, we hypothesize an increase in internal phase volume will produce a polyHIPE foam with a higher porosity, ultimately yielding a softer foam.^32^ Conversely, a decrease in internal phase volume will result in a less porous material, generating a stiffer foam. This highlights the rapid tunability of emulsion templating as a material fabrication platform for designing highly porous materials with target mechanical properties. To build off the information acquired from the previous study, we chose to evaluate the mechanical properties of *star*-PCL-TA foams fabricated from emulsions with internal phases volumes of 70, 75, and 80%. PolyHIPEs are defined as polymerized materials deriving from emulsions containing >74% internal phase volume, determined from the maximum packing capacity of monodispersed spheres.^32^ Alternatively, internal phase volumes from 30-74% result in emulsions that form polymerized medium internal phase emulsions (polyMIPEs) and generally display thicker pore walls and a lower degree of interconnectedness when controlling for all other input variables (e.g. surfactant concentration, mixing speed, etc.). SEM images reveal a visual difference in pore wall thickness and the frequency of pore interconnects (**Figure 4a**). PolyMIPEs displayed incidences of closed pore morphology, whereas polyHIPEs with 80% internal phase demonstrated increased interconnectedness. The compressive modulus of all compositions was determined to be <1 MPa at T = 37°C (hydrated), with increasing foam porosity corresponding with a decrease in modulus. The rapid tunability of these parameters through emulsion inputs is beneficial towards our goal of designing a stent that is compatible with soft tissues and comfortable for the patient throughout the healing period. Although it would be convenient to be able to predict success of the material at maintaining the vaginal lumen from this information, the 3D geometry of the full-size stent (i.e. wall thickness, diameter) will also have a significant impact on its mechanical robustness, therefore convoluting any direct conclusions that may be drawn. This further emphasizes the importance and rationale behind the design of in vitro and in silico models to elucidate the impacts of the vaginal mechanics on gynecological devices.

**Figure 4:**
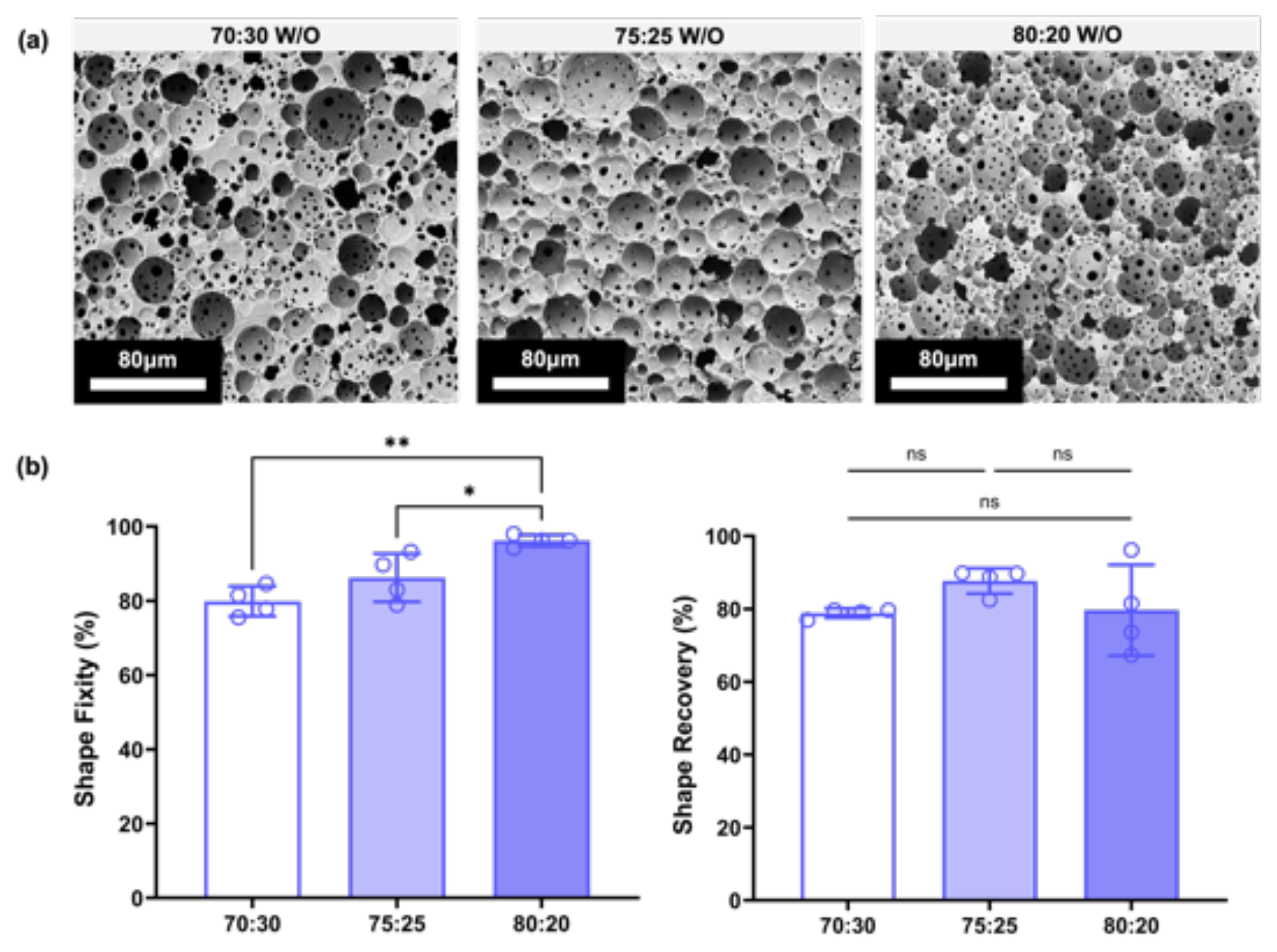
Structural properties of *star*-PCL-TA polyHIPEs with increasing internal phase volumes. (a) Representative SEM images of *star-*PCL-TA foams with increasing internal phase volumes. (b) Shape memory behavior of foams with increasing internal phase volumes, including shape fixity and shape recovery values (T = 37°C, hydrated, n = 4). * = *p* < 0.05, ** = *p* < 0.01, ns = no significance.

The shape memory behavior of *star*-PCL-TA foams with varying internal phase volumes were evaluated to understand the potential impacts of foam porosity and microarchitecture on fixity and recovery (**Figure 4b**). Foams exhibited a statistically significant increase in shape fixity with increasing internal phase volume. Interestingly, the shape recovery results did not mirror this trend, with no significant difference found between their shape recovery. There are multiple factors that may contribute to this, namely the presence of closed pores that restrict trapped air from escaping during compression and insufficient melting of the crystalline lamellae. We hypothesize that higher frequency of closed cell pores within the polyMIPE foams restrict the compressibility of the foam, resulting in plastic deformation to the polymer network that negatively impacts shape fixity and shape recovery of the material. Additionally, internal phase volumes greater than 75% result in a polymer network that lack sufficient density of crystalline lamellae to robustly drive the recovery of its permanent shape at 37°C. From this information, we decided to move forward with an internal phase volume of 75% for full size stent fabrication to ultimately provide the most consistent shape memory behavior to the final device.

**Table 2.**
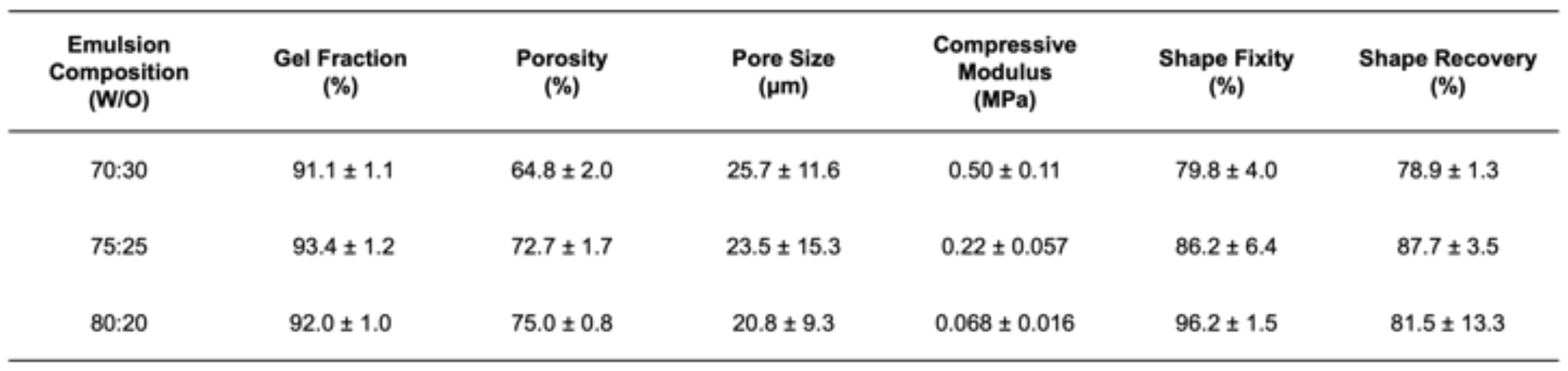
Structural properties of 70:30, 75:25, and 80:20 star-PCL-TA emulsion templated foams.

### 3.3 Vaginal Stent Shape Memory Behavior

Implementation of a custom mold and curing system for the fabrication of SMP polyHIPE vaginal stents successfully yielded hollow, cylindrical stents (**Figure 5a**). The shrinkage that occurred during the polymer network densification resulted in ∼33% decrease in size from the initial fabrication dimensions, resulting in final stents compatible with physiological vaginal dimensions.^31^ Following exposure to *T*_*crimp*_, the stent was crimped to its temporary insertion shape (50% reduction in diameter, *OD* ∼ 11 mm) with a custom radial crimper (**Figure 5b**). The crimping device was designed using concentric 3D printed gear components (**Figure S6**) mounted firmly to a wooden base, modeled after industrial crimpers used for cardiac implants. The protocol used to evaluate radial shape memory was modified to improve upon results from axial shape memory studies, including an increased fixation time, drying *in vacuo*, and increasing the temperature of the water bath used to elicit shape recovery (T = 37°C → T = 45°C). We hypothesized that increasing fixation time in conjunction with vacuum drying would allow more time for crystallization that is responsible for maintaining the temporary shape and removing residual water that may impede lamellar packing. Additionally, increasing the temperature for recovery to 45°C would improve shape recovery by surpassing the *T*_*m*_ of the crystalline regions and completely melting the switching segments of the shape memory system while remaining tissue-safe. The polyHIPE vaginal stents exhibited excellent shape fixity (*R*_*f*_ > 95%) and shape recovery (*R*_*r*_ ∼ 100%) with the revised shape memory protocol (**Figure 5c**). This information was useful in the development of stent deployment protocols for the benchtop testing apparatus, including irrigation with 45°C water to trigger the recovery of the full stent dimensions.

**Figure 5:**
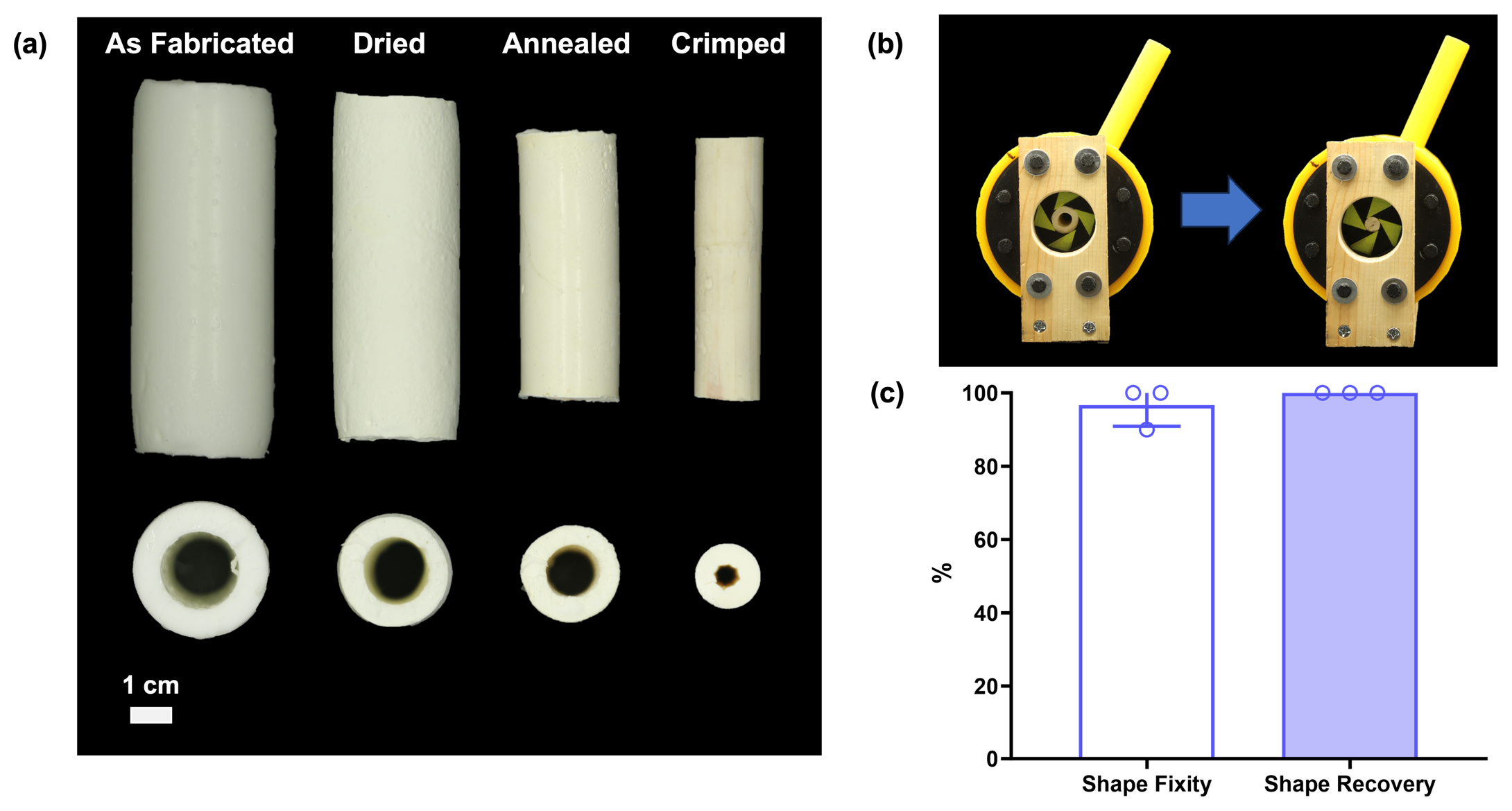
*Star*-PCL-TA polyHIPE (75:25 W/O) vaginal stent. (a) Cross-section images of stents after crimping and re-expansion. (b) Custom crimping device used to radially compress SMP stents into their temporary compressed geometry for ease of deployment. (c) Radial shape memory behavior of full-size stents (T = 45°C, hydrated, n = 3).

### 3.4 Modeling of Vaginal Conditions *In Vitro* and *In Silico*

The benchtop testing apparatus successfully reproduced the physical anatomy, temperatures, and pressures of a vaginal canal (**Figure 6, Figure S7**). Temperature readings taken at three locations within the model (cervical, central, and introitus regions) were confirmed to be within physiological ranges of the vaginal canal (36°C ± 1°C).^33, 34^ Pressures measured by the MizCure perineometer were within the range of PFM contraction pressures reported clinically using this device (19 mmHg ± 1 mmHg).^26^ Additionally, a resting pressure of 10 mmHg was calibrated and used to evaluate the effect of lower pressures on SMP stent deformation. There are three relevant types of forces exerted on the vaginal canal to consider in the design of an *in vitro* simulated pelvic model: resting forces, Valsalva forces, and PFM contractions.^35^ Some clinical data has been acquired and reported with the use of vaginal tactile imaging (VTI), which utilizes a probe equipped with an array of pressure sensors to measure time-varying pressures experienced in the vaginal canal under various conditions.^35-37^ However, one limitation of our benchtop testing apparatus is that it lacks muscle specificity and directionality of those forces by utilizing a constant air pressure to simulate contractions. Therefore, it is more relevant to evaluate resulting *in vitro* pressures as scalar measurements from more wholistic, readily available equipment such as perineometers.

**Figure 6:**
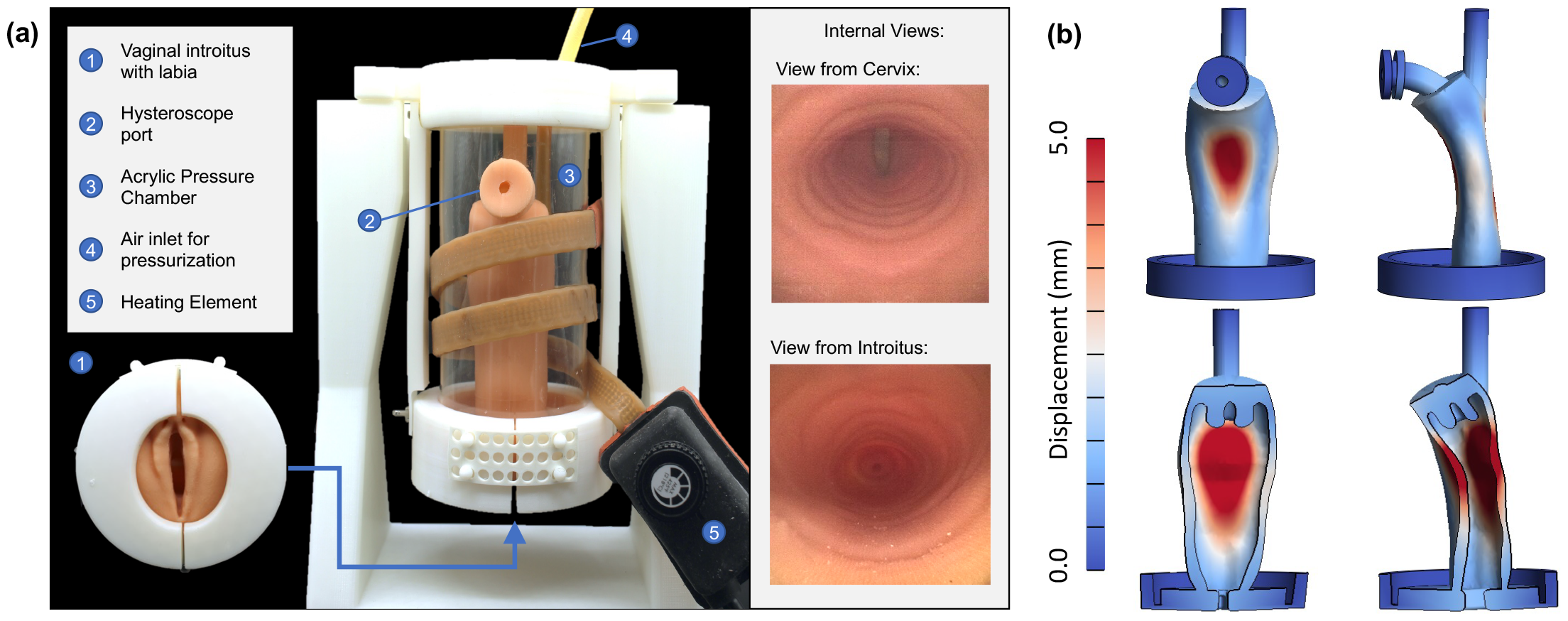
Benchtop testing apparatus used to evaluate in vitro stent deployment and retention. (a) Schematic of benchtop testing apparatus. (b) Complete (top) and cross-sectional (bottom) view of the vaginal wall model under external pressure.

**Figure 7:**
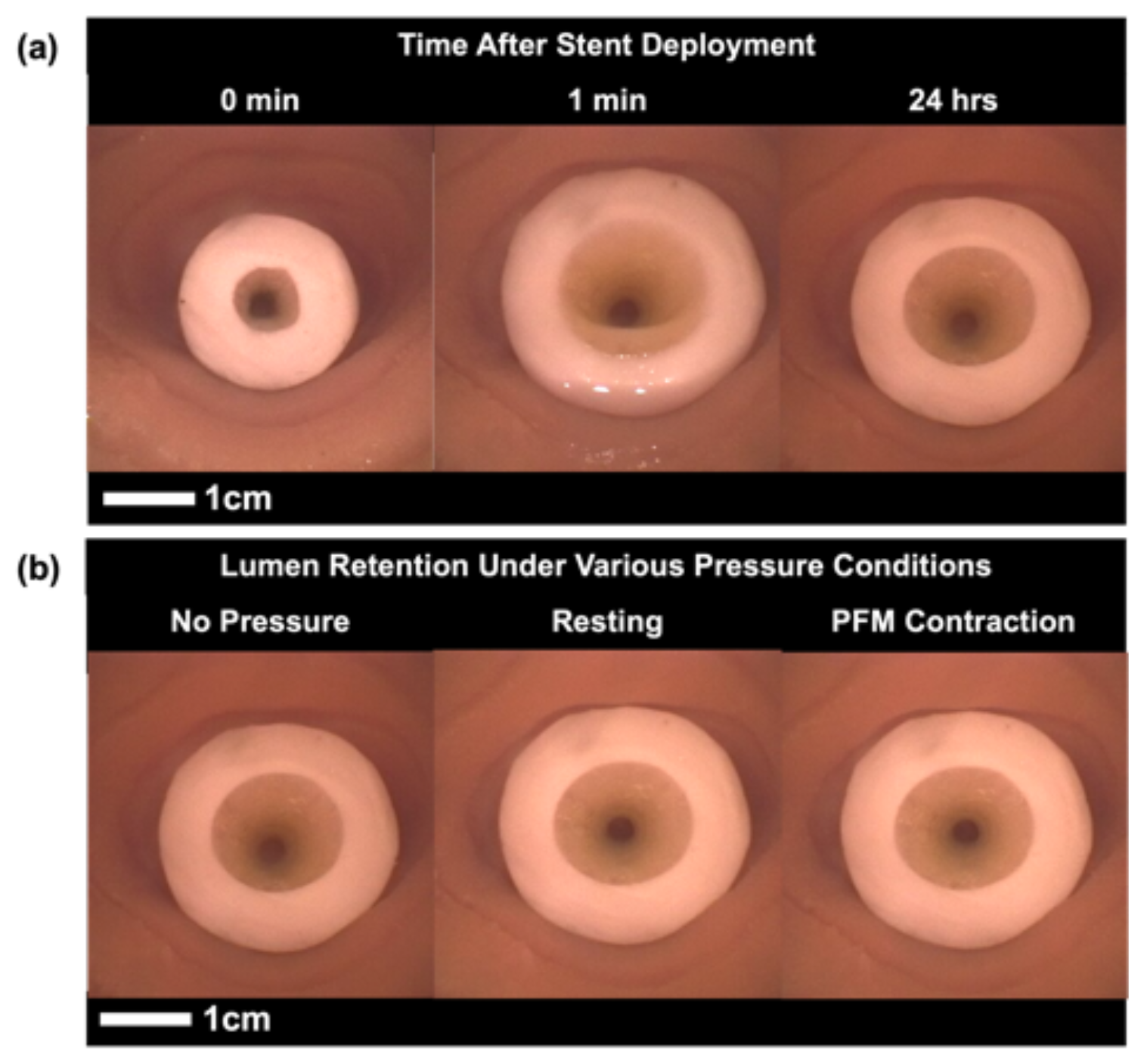
Hysteroscope images of self-fitting stent during expansion and under various pressure conditions. (a) Representative images of stent expansion prior to irrigation, after 1 min, & after 24 hr. (b) Representative images of lumen retention while exposed to resting forces and pelvic floor muscle (PFM) contractions.

The virtual benchtop model successfully executed and predicted the deformation to external pressures while accurately reflecting the imposed boundary conditions. No obvious locking effects were observed while enforcing material incompressibility. Contact between the walls (and between internal devices) was confidently enforced using FEBio’s sliding contact formulation. On a standard laptop computer (Apple M1 Pro), execution wall times were approximately 3-5 minutes, depending on the magnitude of the external pressure. Given the short execution times, this model will be a useful addition to the physical benchtop model and allow for rapid and inexpensive model-directed (virtual) designs of intra-vaginal devices, for example. Future improvements may involve calibration of frictional contact parameters to allow for better prediction of vaginal wall and device interactions, e.g., for device retention studies.

### 3.5 Deployment and Retention of SMP Vaginal Stent *In Vitro*

We elected to evaluate the performance of SMP vaginal stents in response to resting pressures and PFM contractions due to the limited availability of clinical data measured with the MizCure perineometer during the Valsalva maneuver. The largest difference between the forces exerted on the vaginal lumen during a Valsalva maneuver and a voluntary PFM contraction is the directionality and the muscle groups involved in directing the motion.^38^ According to clinical data recorded using VTI, the average pressure generated during voluntary contractions are considerably greater than those experienced during Valsalva maneuvers.^36^ Therefore, it is reasonable to assume that successful performance (which we have defined as <10% lumen deformation for the SMP vaginal stents) under simulated PFM contractions would indicate similar performance under Valsalva pressures.

A crimped SMP polyHIPE vaginal stent expanded to walls of the canal (∼70% increase in cross-sectional area) in <5 minutes after irrigation with warm water (∼45°C). After allowing the stent to equilibrate to physiological temperature overnight, resting and PFM pressures were applied to the stent sequentially and imaged with an Endosee. The stents’ average cross-sectional area decreased by <10% (8.3 ± 7.0%, n = 3) in response to these physiological pressures. Stent diameter exhibited ∼1% (1.1 ± 7.5%, n = 3) decrease along the anterior-posterior dimension, with a corresponding 11% (11.0 ± 3.6%, n = 3) increase distally. Additionally, there was no visible lateral movement of the stent along the vaginal canal due to the simulated physiological pressures. Although this is promising, it is important to note that a limitation of this model is differences in lubricity compared to *in vivo*, which would impact the propensity for a stent to egress from the canal. Overall, this functional evaluation using our novel *in vitro* benchtop testing apparatus indicates an initial proof of concept for a robust, conformable vaginal stent from an emulsion-templated SMP polyHIPE.

## 4. Conclusion

In this work, we report the iterative development of a SMP vaginal stent, from material design to functional testing modalities *in vitro* and *in silico*. First, we synthesized a new PCL photocurable macromer that could be fabricated into a SMP foam with rapid shape recovery at physiological temperature. We also evaluated emulsion templating fabrication inputs to modify bulk mechanical properties of resulting SMP foams. By implementing fundamental understandings of known structure-property relationships on both a molecular and microarchitectural level, we successfully integrated our carefully engineered material in the design of a full-size vaginal stent. Our device exhibits the ability to maintain a temporary shape for ease of insertion into the vaginal canal and expands upon application of warm water to the walls of the canal to anatomically secure its position, limiting stent egress. In addition to providing evidence of successful stent deployment and retention, the *in vitro* and *in silico* pelvic models described provide an accessible framework for the iterative design of future gynecological devices.

