## Supplemental Figures for "Polycaprolactone-based shape memory foams as self-fitting vaginal stents"

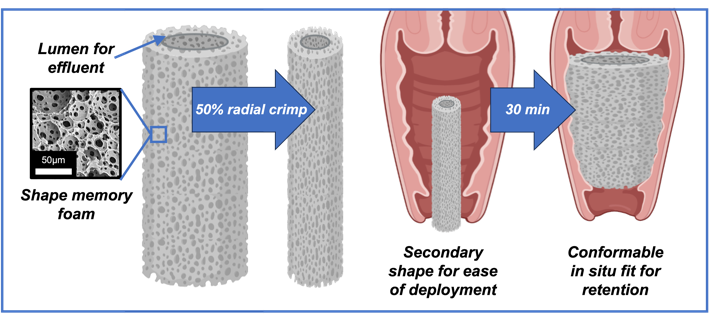

Created in BioRender

1. **Introduction**

There is an urgent clinical need for a vaginal stent that can address vaginal stenosis in post-surgical and post-radiation settings, impacting over 340,000 patients in North America each year.^1^^,^ ^2^ The urgency in addressing vaginal stenosis is underscored by the shortcomings of current treatment options. Existing commercially available stents, including stainless steel variants and inflatable balloon stents, exhibit notable challenges with comfort and retention in the vagina and often require additional fixation to maintain their position in the canal. Stent expulsion can occur during simple routine motions like walking or sitting, making them highly ineffective solutions for most patients. As a result, physicians are left to craft makeshift stents from common medical equipment such as foley catheters, condoms, and medical foam.^3^^,^ ^4^ This is particularly true in patients whose size is incompatible with current commercially available options, most often pediatric patients. The search for an alternative solution remains underdeveloped, resulting in ~50% rate of secondary revision surgery for pediatric patients.^5^ The failure modes of these devices indicate that the design of a novel vaginal stent must rely on the premise of a conformal, retained fit within the vaginal cavity.

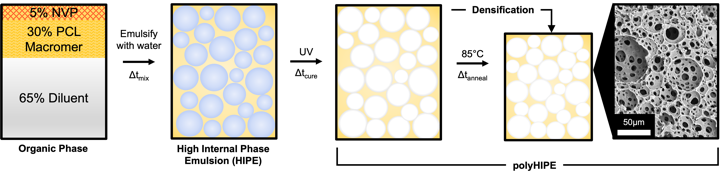

**Figure 1**: General scheme of emulsion templating of shape memory foams. The internal phase is gradually added into the organic phase while the system is mixed, forming a high internal phase emulsion (HIPE). The material is then cured to form a polymerized high internal phase emulsion (polyHIPE).

2.4 **Gel Fraction**

The gel fraction was measured gravimetrically to evaluate the extent of network formation. First, samples (*d* ~ 6 mm, *h* ~ 1 mm, n = 5) were weighed after drying *in vacuo* for 48 hours. The samples were placed in dichloromethane (DCM) for 4 hours at 1 mL / 10 mg of the sample, and vacuum dried again until a constant mass was achieved. The final weight divided by the initial weight was assessed as the gel fraction and corrected for the mass of the surfactant that would dissolve in the DCM.

$$Gel fraction (\%)=\left( \frac{Final Mass}{Corrected Mass} \right)*100$$

***Equation 1****: Gel Fraction*

2.5 **Porosity**

The porosity of PCL polyHIPEs was determined gravimetrically. First, the polymer density was measured from solid film specimens of each macromer (*d* ~ 6 mm, *h* ~ 2 mm, n = 5). Dried HIPE samples (*d* ~ 6 mm, *h* ~ 2 mm, n = 5) were measured with electronic calipers and weighed. Porosity was calculated as a percent difference in the density of foams and analogous films.

$$R_{f}\left( N \right)= \frac{\varepsilon_{u}(N)}{\varepsilon_{m}}$$

***Equation 3****: Shape Fixity*

$$R_{r}\left( N \right)= \frac{\varepsilon_{m}-\varepsilon_{p}(N)}{\varepsilon_{m}-\varepsilon_{p}(N-1)}$$

***Equation 4****: Shape Recovery*

2.9 **Compressive Properties**

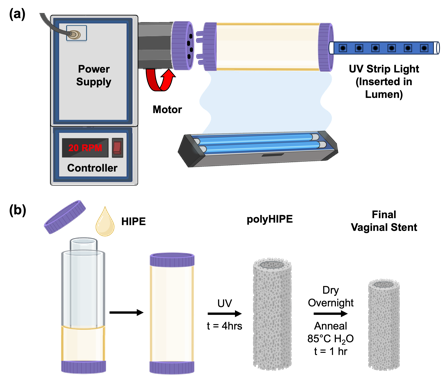

**Figure 2**: Fabrication of PCLTA polyHIPE vaginal stents. (a) Custom rotational device for UV curing of the vaginal stent. (b) General scheme of stent fabrication demonstrating mold setup and densification after fabrication. Created in BioRender.

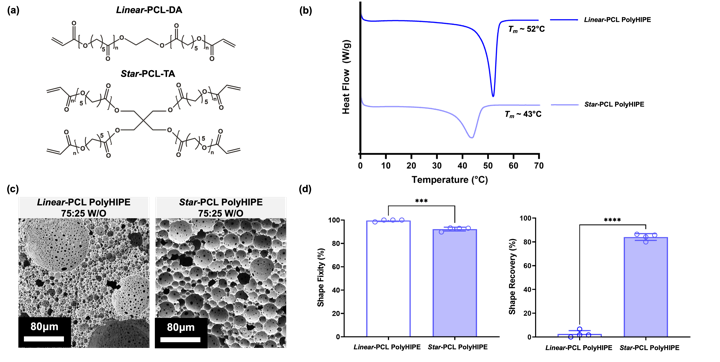

**Figure 3**: Structural properties of synthesized *linear*-PCL-DA and *star*-PCL-TA polyHIPE foams. (a) Schematic representing the chemical structures of the *linear*-PCL-DA and *star*-PCL-TA. (b) DSC thermograms of *linear* and *star* foam compositions demonstrating a decrease in *T_m_* with *star* architecture. (c) Representative SEM images of *linear* and *star* foams. (d) Shape memory behavior of *linear* and *star* foams, including shape fixity and shape recovery values (T = 37°C, hydrated, n = 4). *** = p < 0.001, **** = p < 0.0001.

**Table 1**. Structural properties of linear-PCL-DA and star-PCL-TA polyHIPEs (75:25 W/O).

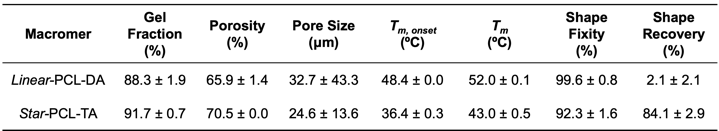

3.2 **Effect of Emulsion Internal Phase Volume on SMP PolyHIPE Properties**

The effect of the SMP HIPE’s internal phase volume on resulting foam porosity was used to influence the *star*-PCL-TA polyHIPE’s mechanical properties. Based on the principles of emulsion templating, we hypothesize an increase in internal phase volume will produce a polyHIPE foam with a higher porosity, ultimately yielding a softer foam.^32^ Conversely, a decrease in internal phase volume will result in a less porous material, generating a stiffer foam. This highlights the rapid tunability of emulsion templating as a material fabrication platform for designing highly porous materials with target mechanical properties. To build off the information acquired from the previous study, we chose to evaluate the mechanical properties of *star*-PCL-TA foams fabricated from emulsions with internal phases volumes of 70, 75, and 80%. PolyHIPEs are defined as polymerized materials deriving from emulsions containing >74% internal phase volume, determined from the maximum packing capacity of monodispersed spheres.^32^ Alternatively, internal phase volumes from 30-74% result in emulsions that form polymerized medium internal phase emulsions (polyMIPEs) and generally display thicker pore walls and a lower degree of interconnectedness when controlling for all other input variables (e.g. surfactant concentration, mixing speed, etc.). SEM images reveal a visual difference in pore wall thickness and the frequency of pore interconnects (**Figure 4a**). PolyMIPEs displayed incidences of closed pore morphology, whereas polyHIPEs with 80% internal phase demonstrated increased interconnectedness. The compressive modulus of all compositions was determined to be <1 MPa at T = 37°C (hydrated), with increasing foam porosity corresponding with a decrease in modulus. The rapid tunability of these parameters through emulsion inputs is beneficial towards our goal of designing a stent that is compatible with soft tissues and comfortable for the patient throughout the healing period. Although it would be convenient to be able to predict success of the material at maintaining the vaginal lumen from this information, the 3D geometry of the full-size stent (i.e. wall thickness, diameter) will also have a significant impact on its mechanical robustness, therefore convoluting any direct conclusions that may be drawn. This further emphasizes the importance and rationale behind the design of in vitro and in silico models to elucidate the impacts of the vaginal mechanics on gynecological devices.

The shape memory behavior of *star*-PCL-TA foams with varying internal phase volumes were evaluated to understand the potential impacts of foam porosity and microarchitecture on fixity and recovery (**Figure 4b**). Foams exhibited a statistically significant increase in shape fixity with increasing internal phase volume. Interestingly, the shape recovery results did not mirror this trend, with no significant difference found between their shape recovery. There are multiple factors that may contribute to this, namely the presence of closed pores that restrict trapped air from escaping during compression and insufficient melting of the crystalline lamellae. We hypothesize that higher frequency of closed cell pores within the polyMIPE foams restrict the compressibility of the foam, resulting in plastic deformation to the polymer network that negatively impacts shape fixity and shape recovery of the material. Additionally, internal phase volumes greater than 75% result in a polymer network that lack sufficient density of crystalline lamellae to robustly drive the recovery of its permanent shape at 37°C. From this information, we decided to move forward with an internal phase volume of 75% for full size stent fabrication to ultimately provide the most consistent shape memory behavior to the final device.

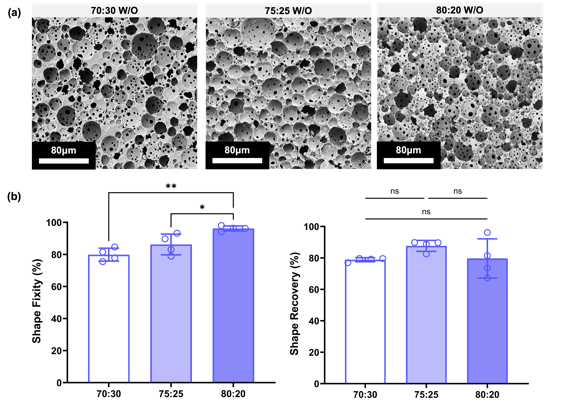

**Figure 4**: Structural properties of *star*-PCL-TA polyHIPEs with increasing internal phase volumes. (a) Representative SEM images of *star-*PCL-TA foams with increasing internal phase volumes. (b) Shape memory behavior of foams with increasing internal phase volumes, including shape fixity and shape recovery values (T = 37°C, hydrated, n = 4). * = *p* < 0.05, ** = *p* < 0.01, ns = no significance.

**Table 2**. Structural properties of 70:30, 75:25, and 80:20 star-PCL-TA emulsion templated foams.

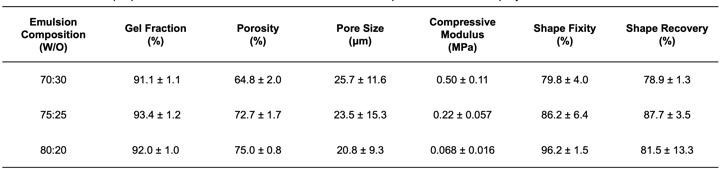

3.3 **Vaginal Stent Shape Memory Behavior**

Implementation of a custom mold and curing system for the fabrication of SMP polyHIPE vaginal stents successfully yielded hollow, cylindrical stents (**Figure 5a**). The shrinkage that occurred during the polymer network densification resulted in ~33% decrease in size from the initial fabrication dimensions, resulting in final stents compatible with physiological vaginal dimensions.^31^ Following exposure to *T_crimp_*, the stent was crimped to its temporary insertion shape (50% reduction in diameter, *OD* ~ 11 mm) with a custom radial crimper (**Figure 5b**). The crimping device was designed using concentric 3D printed gear components (**Figure S6**) mounted firmly to a wooden base, modeled after industrial crimpers used for cardiac implants. The protocol used to evaluate radial shape memory was modified to improve upon results from axial shape memory studies, including an increased fixation time, drying *in vacuo*, and increasing the temperature of the water bath used to elicit shape recovery (T = 37°C → T = 45°C). We hypothesized that increasing fixation time in conjunction with vacuum drying would allow more time for crystallization that is responsible for maintaining the temporary shape and removing residual water that may impede lamellar packing. Additionally, increasing the temperature for recovery to 45°C would improve shape recovery by surpassing the *T_m_* of the crystalline regions and completely melting the switching segments of the shape memory system while remaining tissue-safe. The polyHIPE vaginal stents exhibited excellent shape fixity (*R_f_* > 95%) and shape recovery (*R_r_* ~ 100%) with the revised shape memory protocol (**Figure 5c**). This information was useful in the development of stent deployment protocols for the benchtop testing apparatus, including irrigation with 45°C water to trigger the recovery of the full stent dimensions.

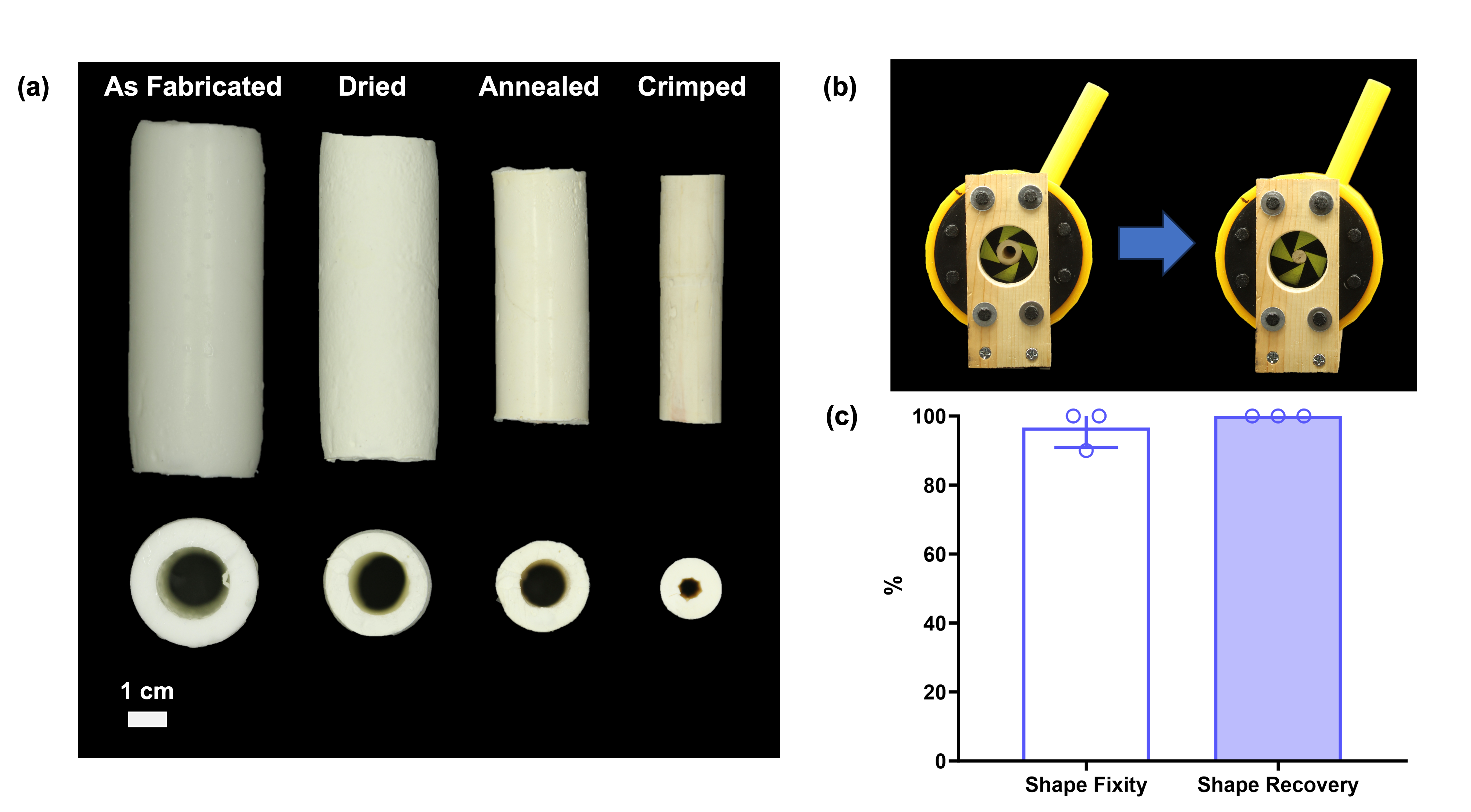

**Figure 5**: *Star*-PCL-TA polyHIPE (75:25 W/O) vaginal stent. (a) Cross-section images of stents after crimping and re-expansion. (b) Custom crimping device used to radially compress SMP stents into their temporary compressed geometry for ease of deployment. (c) Radial shape memory behavior of full-size stents (T = 45°C, hydrated, n = 3).

3.4 **Modeling of Vaginal Conditions *In Vitro* and *In Silico***

The benchtop testing apparatus successfully reproduced the physical anatomy, temperatures, and pressures of a vaginal canal (**Figure 6, Figure S7**). Temperature readings taken at three locations within the model (cervical, central, and introitus regions) were confirmed to be within physiological ranges of the vaginal canal (36°C ± 1°C).^33^^,^ ^34^ Pressures measured by the MizCure perineometer were within the range of PFM contraction pressures reported clinically using this device (19 mmHg ± 1 mmHg).^26^ Additionally, a resting pressure of 10 mmHg was calibrated and used to evaluate the effect of lower pressures on SMP stent deformation. There are three relevant types of forces exerted on the vaginal canal to consider in the design of an *in vitro* simulated pelvic model: resting forces, Valsalva forces, and PFM contractions.^35^ Some clinical data has been acquired and reported with the use of vaginal tactile imaging (VTI), which utilizes a probe equipped with an array of pressure sensors to measure time-varying pressures experienced in the vaginal canal under various conditions.^35-37^ However, one limitation of our benchtop testing apparatus is that it lacks muscle specificity and directionality of those forces by utilizing a constant air pressure to simulate contractions. Therefore, it is more relevant to evaluate resulting *in vitro* pressures as scalar measurements from more wholistic, readily available equipment such as perineometers.

The virtual benchtop model successfully executed and predicted the deformation to external pressures while accurately reflecting the imposed boundary conditions. No obvious locking effects were observed while enforcing material incompressibility. Contact between the walls (and between internal devices) was confidently enforced using FEBio’s sliding contact formulation. On a standard laptop computer (Apple M1 Pro), execution wall times were approximately 3-5 minutes, depending on the magnitude of the external pressure. Given the short execution times, this model will be a useful addition to the physical benchtop model and allow for rapid and inexpensive model-directed (virtual) designs of intra-vaginal devices, for example. Future improvements may involve calibration of frictional contact parameters to allow for better prediction of vaginal wall and device interactions, e.g., for device retention studies.
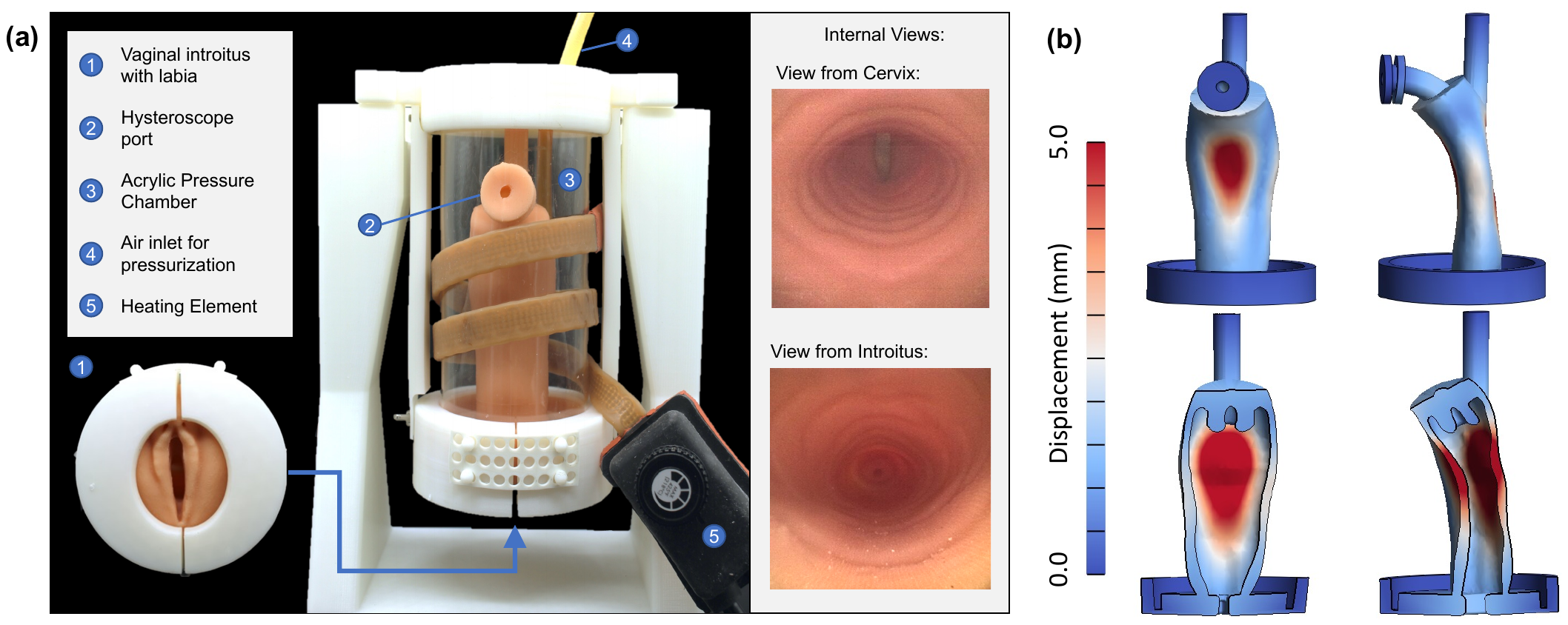

**Figure 6**: Benchtop testing apparatus used to evaluate in vitro stent deployment and retention. (a) Schematic of benchtop testing apparatus. (b) Complete (top) and cross-sectional (bottom) view of the vaginal wall model under external pressure.

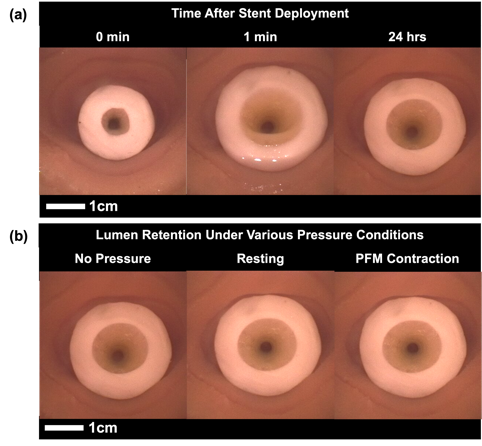

**Figure 7**: Hysteroscope images of self-fitting stent during expansion and under various pressure conditions. (a) Representative images of stent expansion prior to irrigation, after 1 min, & after 24 hr. (b) Representative images of lumen retention while exposed to resting forces and pelvic floor muscle (PFM) contractions.

4. **Conclusion**

In this work, we report the iterative development of a SMP vaginal stent, from material design to functional testing modalities *in vitro* and *in silico*. First, we synthesized a new PCL photocurable macromer that could be fabricated into a SMP foam with rapid shape recovery at physiological temperature. We also evaluated emulsion templating fabrication inputs to modify bulk mechanical properties of resulting SMP foams. By implementing fundamental understandings of known structure-property relationships on both a molecular and microarchitectural level, we successfully integrated our carefully engineered material in the design of a full-size vaginal stent. Our device exhibits the ability to maintain a temporary shape for ease of insertion into the vaginal canal and expands upon application of warm water to the walls of the canal to anatomically secure its position, limiting stent egress. In addition to providing evidence of successful stent deployment and retention, the *in vitro* and *in silico* pelvic models described provide an accessible framework for the iterative design of future gynecological devices.

**Supplemental Figures:**

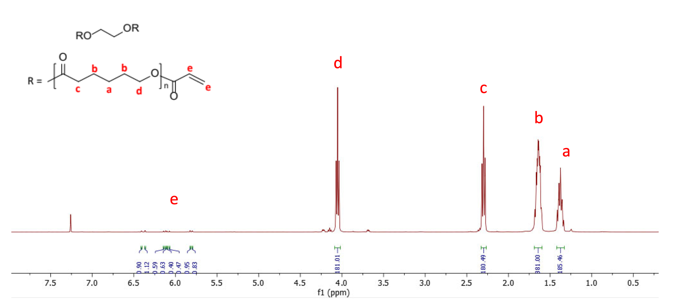

**(a)** ^1^H NMR of “***n = 90***” Linear‐PCL‐DA, (δ, ppm): **a**: 1.33‐1.42 (180H, OCHCH_2_CH_2_***CH_2_***CH_2_CH_2_O), **b**: 1.59‐1.69 (360H, OCHCH_2_***CH_2_***CH_2_***CH_2_***CH_2_O) **c**: 2.27‐2.33 (180H, OCH***CH_2_***CH_2_CH_2_CH_2_CH_2_O), **d**: 4.02‐4.08 (180H, OCHCH_2_CH_2_CH_2_CH_2_***CH_2_***O), **e**: 5.79‐5.81, 5.82‐5.83, 6.07‐6.08, 6.09‐6.10, 6.11‐6.12, 6.13‐6.15, 6.36‐6.37, 6.40‐6.42 (6H, OC***CHCH_2_***)

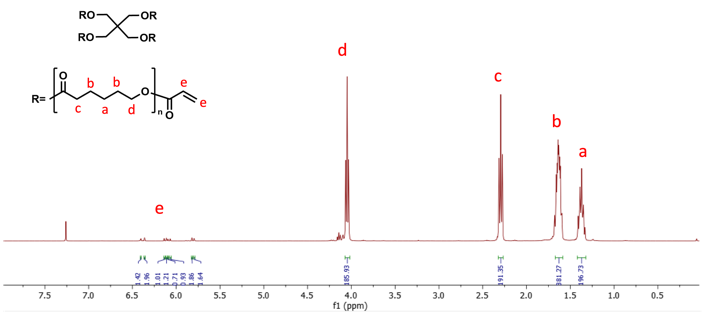

**(b)** ^1^H NMR of “***n = 88***” Star-PCL-TA, (δ, ppm): **a**: 1.30-1.39 (176H, OCHCH_2_CH_2_***CH_2_***CH_2_CH_2_O), **b**: 1.56-1.66 (352H, OCHCH_2_***CH_2_***CH_2_***CH_2_***CH_2_O) **c**: 2.24-2.30 (176H, OCH***CH_2_***CH_2_CH_2_CH_2_CH_2_O), **d**: 3.99-4.05 (176H, OCHCH_2_CH_2_CH_2_CH_2_***CH_2_***O), **e**: 5.76-5.77, 5.79-5.80, 6.03-6.05, 6.06-6.07, 6.08-6.09, 6.11-6.12, 6.33-6.34, 6.37-6.38 (12H, OC***CHCH_2_***)

**Figure S1:** Representative NMR Spectra of PCL Macromers. (a) NMR of *linear*-PCL-DA. (b) NMR of *star*-PCL-TA.

**Table S1**. Properties of linear-PCL-DA and star-PCL-TA synthetic batches.

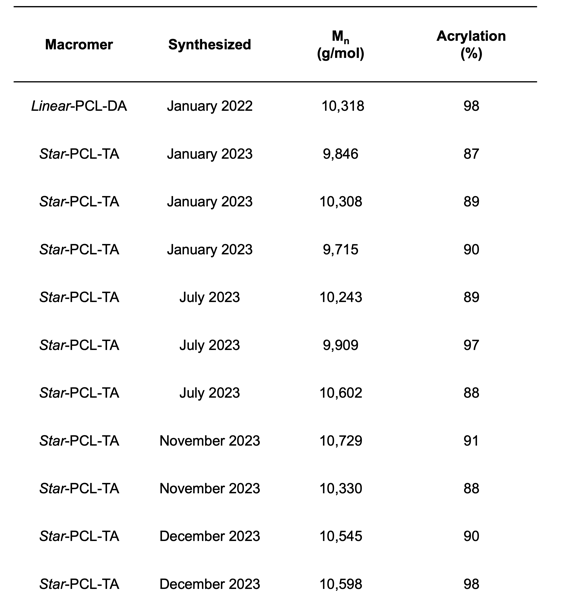

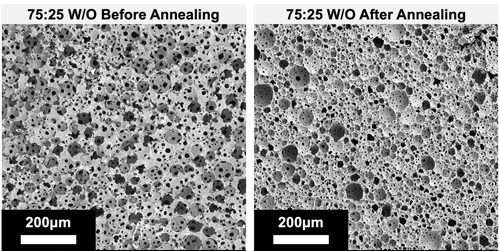

**Figure S2**: SEM images of *star*-PCL-TA polyHIPEs before and after annealing.

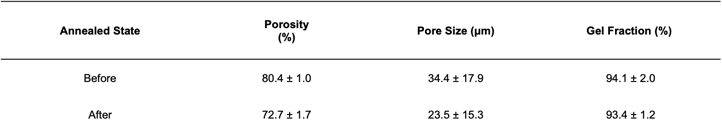
**Table S2**. Structural properties of 75:25 emulsion templated star-PCL-TA polyHIPEs before and after annealing (T = 85°C).

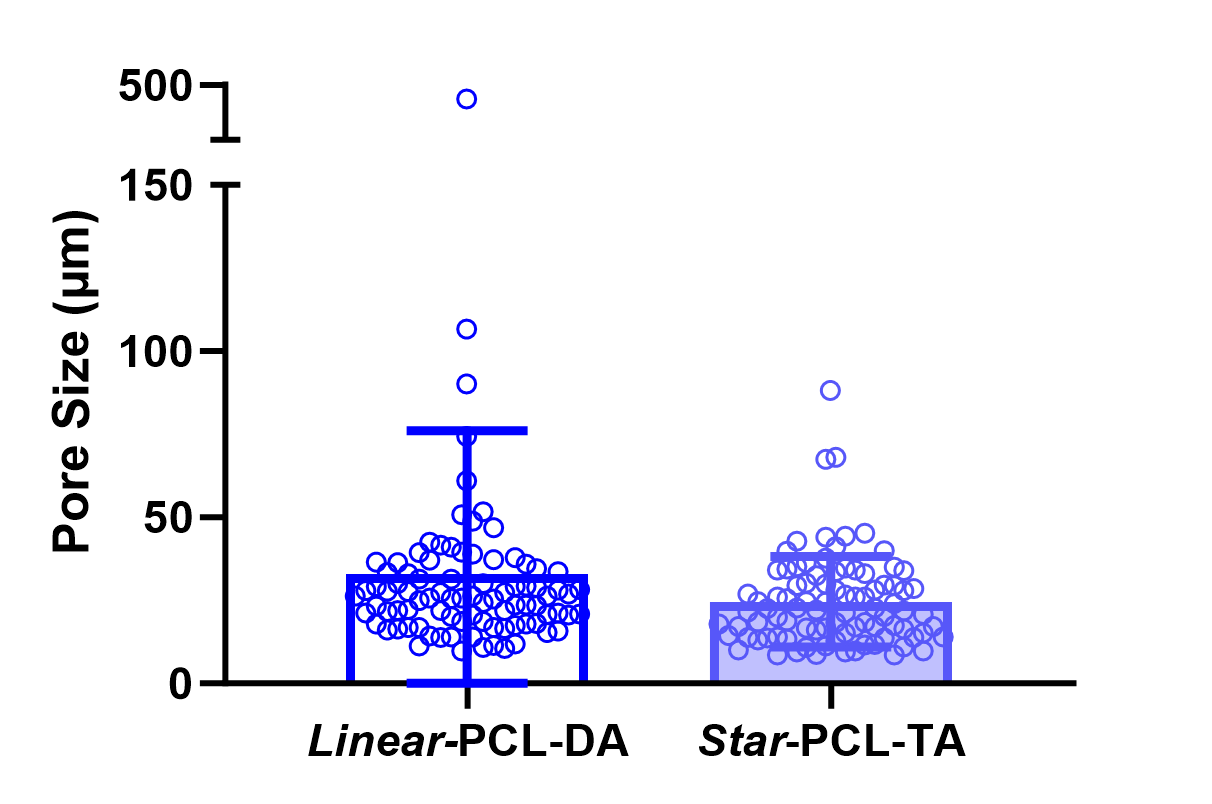

**Figure S3:** Pore Size Distribution of *Linear*-PCL-DA and *star*-PCL-TA polyHIPE foam specimen (n = 9 images).

**Figure S4**: DSC Thermograms of *star*-PCL-TA polyHIPEs at various stages of fabrication and shape memory cycling. (a) First DSC heating cycle (T = 0-200°C, 5°C/min). (b) Second DSC heating cycle (T = 0-200°C, 5°C/min).

**Table S4**: *T_m_* and *T_m,onset_* of 75:25 PCL-TA polyHIPEs at various stages of fabrication and shape memory cycling.

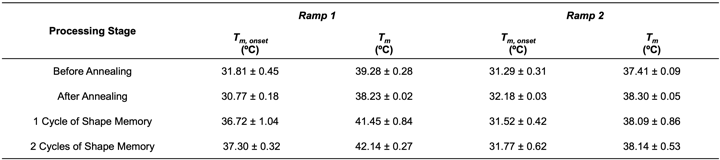

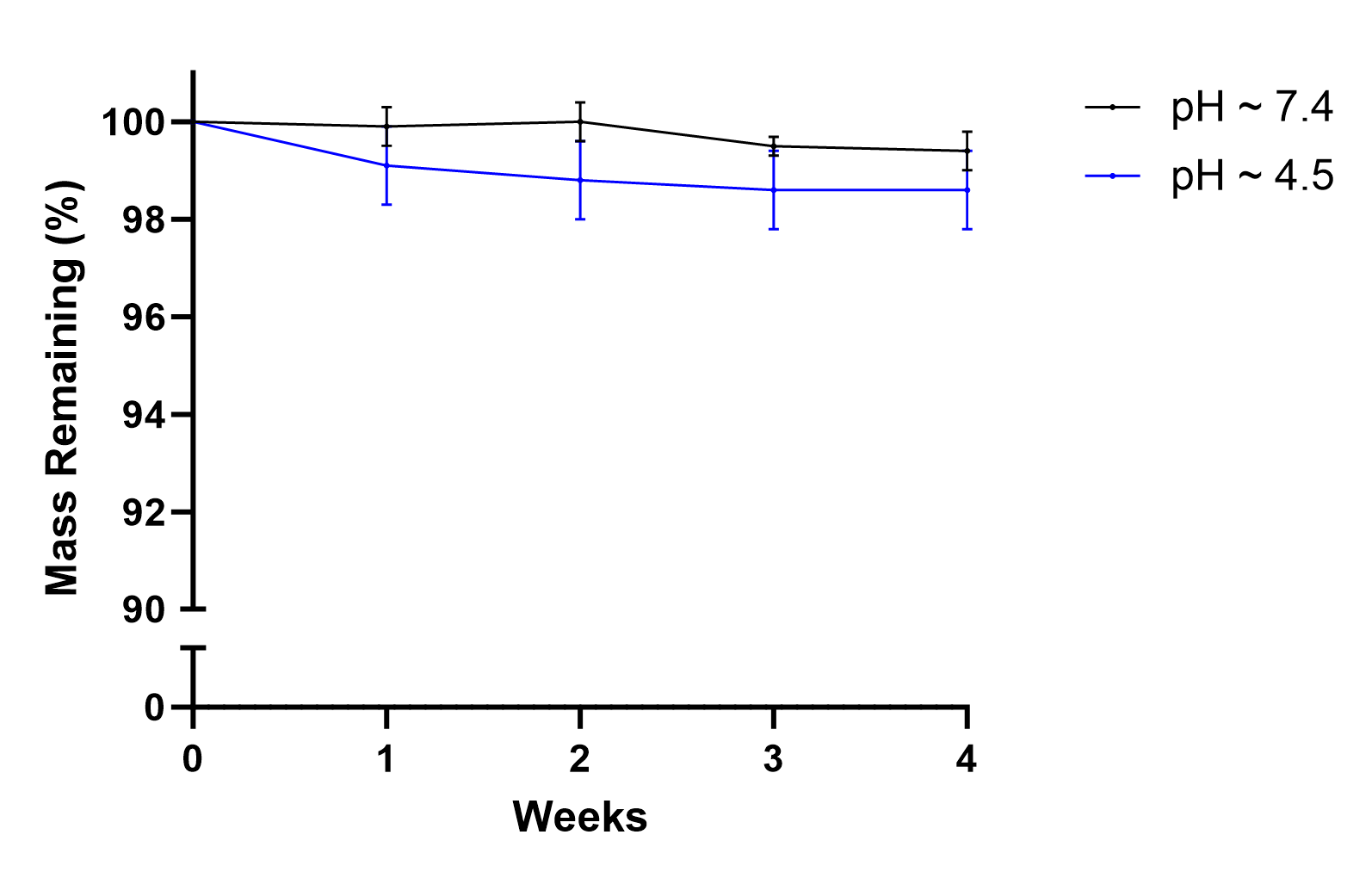

**Figure S5**: Evaluation of real time degradation (t = 4wks) under prepubescent (pH ~7.4) and adolescent/adult conditions (pH ~4.5). Results indicate gravimetric mass loss over time for 75:25 *star*-PCL-TA scaffolds.

**(a)**

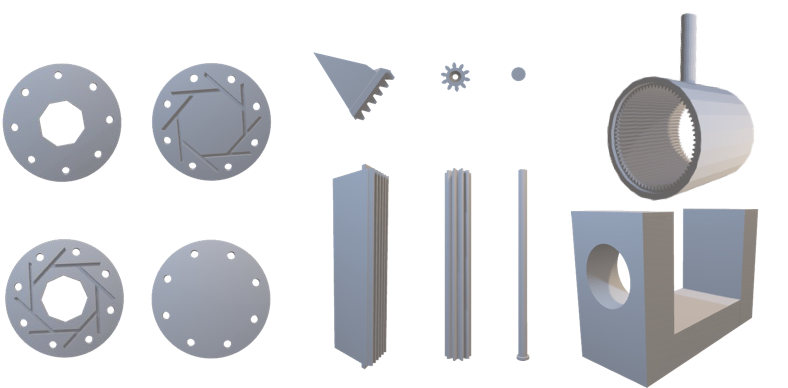

**(b)**

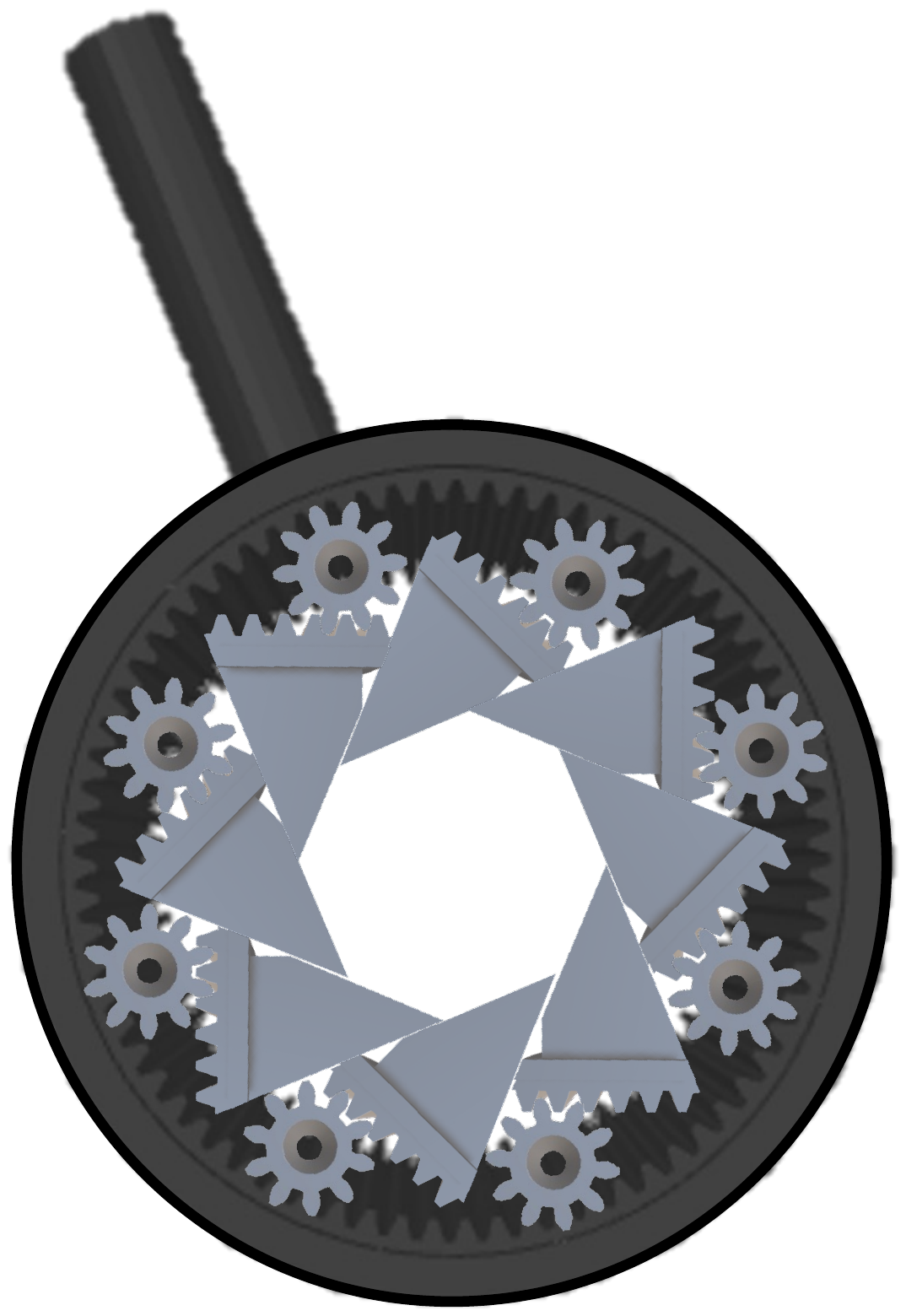

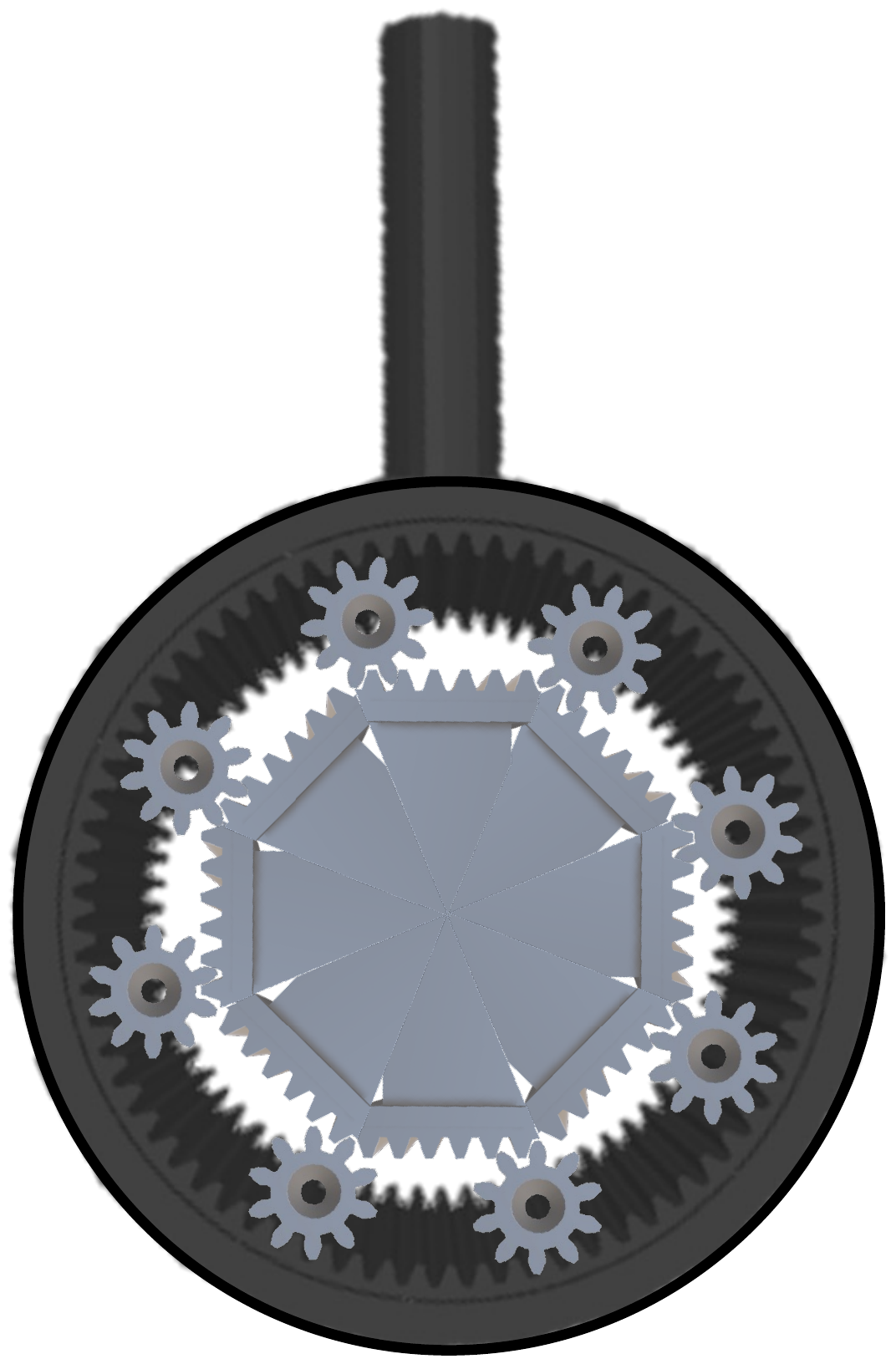

**Figure S6**: Custom radial crimping device. (a) Components of radial crimping device (representative, not to scale). (b) Innerworkings of radial crimping device.

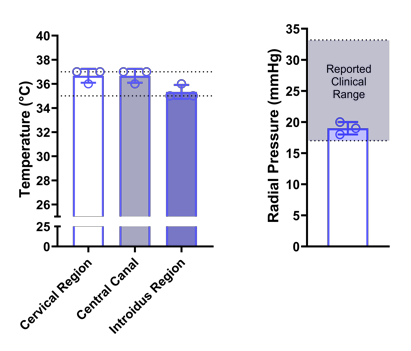

**Figure S7**: Physiological parameters of the benchtop pelvic model, including temperature and pressure measured with a MizCure perineometer.

3.	Tharihalli, C., Muralikrishna V., Shiva Kumar H. C., Surgical management of vaginal agenesis using a modified Mc Indoe’s

technique: VIMS experience. *International Journal of Reproduction, Contraception, Obstetrics and Gynecology* **2017,** *6* (9), 3841-3845.

4.	Fowler, K. G.; Mohindra, P.; Kim, A.; Gomez-Lobo, V., Foley Catheter as a Vaginal Stent in a Toddler with Vaginal Rhabdomyosarcoma. *J Pediatr Adolesc Gynecol* **2018,** *31* (3), 315-317.

5.	Godbole P, K. M., Wilcox D, editors., *Pediatric urology: surgical complications and management*. 2nd ed.; Wiley Blackwell: 2015.

17.	Nail, L. N.; Zhang, D.; Reinhard, J. L.; Grunlan, M. A., Fabrication of a Bioactive, PCL-based "Self-fitting" Shape Memory Polymer Scaffold. *J Vis Exp* **2015,** (105), e52981.

24.	Nail, L. N.; Zhang, D.; Reinhard, J. L.; Grunlan, M. A., Fabrication of a Bioactive, PCL-based "Self-fitting" Shape Memory Polymer Scaffold. *JoVE* **2015,** (104), e52981.

25.	Carnachan, R. J.; Bokhari, M.; Przyborski, S. A.; Cameron, N. R., Tailoring the morphology of emulsion-templated porous polymers. *Soft Matter* **2006,** *2* (7), 608-616.

26.	Abe-Takahashi, Y.; Kitta, T.; Ouchi, M.; Okayauchi, M.; Chiba, H.; Higuchi, M.; Togo, M.; Shinohara, N., Reliability and validity of pelvic floor muscle strength assessment using the MizCure perineometer. *BMC Women's Health* **2020,** *20* (1), 257.

27.	Liao, Z.; Hossain, M.; Yao, X., Ecoflex polymer of different Shore hardnesses: Experimental investigations and constitutive modelling. *Mechanics of Materials* **2020,** *144*, 103366.

28.	Xie, W.; Jiang, N.; Gan, Z., Effects of multi-arm structure on crystallization and biodegradation of star-shaped poly(epsilon-caprolactone). *Macromol Biosci* **2008,** *8* (8), 775-84.

29.	Lu, Y.; An, L.; Wang, Z.-G., Intrinsic Viscosity of Polymers: General Theory Based on a Partially Permeable Sphere Model. *Macromolecules* **2013,** *46* (14), 5731-5740.

30.	Douglas, J. F.; Roovers, J.; Freed, K. F., Characterization of branching architecture through "universal" ratios of polymer solution properties. *Macromolecules* **1990,** *23* (18), 4168-4180.

31.	Barnhart, K. T.; Izquierdo, A.; Pretorius, E. S.; Shera, D. M.; Shabbout, M.; Shaunik, A., Baseline dimensions of the human vagina. *Human Reproduction* **2006,** *21* (6), 1618-1622.

32.	Aldemir Dikici, B.; Claeyssens, F., Basic Principles of Emulsion Templating and Its Use as an Emerging Manufacturing Method of Tissue Engineering Scaffolds. *Front Bioeng Biotechnol* **2020,** *8*, 875.

33.	Boyd, P.; Desjardins, D.; Kumar, S.; Fetherston, S. M.; Le-Grand, R.; Dereuddre-Bosquet, N.; Helgadóttir, B.; Bjarnason, Á.; Narasimhan, M.; Malcolm, R. K., A temperature-monitoring vaginal ring for measuring adherence. *PLoS One* **2015,** *10* (5), e0125682.
